# Microtubule remodeling by the innate immune factor Trim69 compromises dynein-dependent migration of HIV virion cores towards the nucleus

**DOI:** 10.1101/2025.02.06.636796

**Authors:** Charlotte Vadon, Xuan-Nhi Nguyen, Valerie Siahaan, Arya Krishnan, Veronique Henriot, Amandine Chantharath, Julien Burlaud-Gaillard, Philippe Roingeard, Carsten Janke, Lucie Etienne, Francesca Fiorini, Maria M. Magiera, Andrea Cimarelli

## Abstract

Like many viruses, HIV relies on the microtubule (MT) cytoskeleton for successful infection. MT-associated proteins (MAPs) regulate MT functions and thus bear the potential to modulate viral infection. However, while several MAPs are known to exert pro-viral effects on HIV, little is known about antiviral ones. We previously described the Tripartite motif protein 69 (Trim69) as an innate immune factor that remodels MTs, leading to inhibition of a broad spectrum of viruses, including HIV. Through in silico modeling, TIRF microscopy and cell-based assays, we determine that Trim69 binding to MTs is determined by a basic surface in its SPRY domain that interacts with the C-terminal tails of tubulins. This surface is conserved among mammalian Trim69s and is critical for its functions. We demonstrate that by binding and remodeling MTs, Trim69 inhibits the docking and the migration of virion cores on MTs by promoting the stalling of the dynein/dynactin motor complexes. Altogether, these findings shed light on a novel mechanism of viral defense that involves an innate immune regulation of the MT cytoskeleton.

## Introduction

For successful infection, the HIV viral genetic material must integrate into the host genome. For this to occur, the virion cores that are released into the cell cytoplasm upon viral-to-cellular membrane fusion must obligatorily navigate their way into the nucleus, while simultaneously accommodating the process of reverse transcription of the viral RNA genome into double-stranded DNA ^1^.

Previous studies have shown that HIV virion cores entering at the cell periphery hijack the dynein motor complex for retrograde transport along microtubules (MTs) to reach the nucleus ^2^ and several MT-associated proteins (MAPs) have been identified as facilitators of this process ^3-13^. However, little is known about whether cellular defense mechanisms can actively modulate MTs to inhibit viral infection. MTs are polarized tubular structures composed of α/β-tubulin heterodimers that nucleate from a microtubule organizing center (MTOCs) in the proximity of the nucleus. MT minus ends are anchored at the MTOC, while their plus ends point and polymerize towards the cell periphery. MT dynamics and functions in cells are regulated by various parameters including the interaction with a wide spectrum of MAPs and post-translational modifications (PTMs) of tubulins, such as acetylation and detyrosination ^14-16^. While it remains unclear whether these modifications drive MT stabilization or are a consequence of it, their accumulation on stable MTs influences resistance to mechanical stress, MAP-MT interactions, and MT-based transport (reviewed in ^17^).

We recently identified the Tripartite motif protein 69 (Trim69), a member of the large Trim family, as an interferon-induced, innate immune factor capable of binding and remodeling MTs, leading to the inhibition of viruses as diverse as HIV-1, SARS-CoV-2, or the Vesicular Stomatitis RhabdoVirus (VSV) ^18^. This together with results from other studies ^19-22^, establish Trim69 as a *bona fide* broad-spectrum viral inhibitor. Trim69 induces a profound reorganization of the MT network, resulting in increased levels of α-tubulin acetylation and detyrosination, two post-translational modifications typically associated with stable MTs ^18^.

In order to dissect the mechanism of the antiviral activity of Trim69, we combined *in silico* modeling-guided mutagenesis, cell-based assays and *in vitro* reconstitution approaches to characterize how the interaction between Trim69 and MTs affects the intracellular behavior of HIV cores.

Our results indicate that a basic surface in the SPRY domain of Trim69, which is conserved among mammalian Trim69 proteins, mediates its interaction with the acidic C-terminal tails of tubulins. We demonstrate that this interaction is crucial for both MT remodeling and antiviral activities of Trim69. Trim69-dependent MT reorganization disrupts both the docking of HIV virion cores to MTs at the cell periphery and their dynein-dependent transport towards the cell nucleus. Our data show that this is achieved by redistributing and sequestering dynein and dynactin motor complexes onto MTs decorated with Trim69.

Our findings reveal a novel antiviral mechanism involving the reorganization of the MT cytoskeleton, which hinders the efficient transport of viral cores to the nucleus. These results also highlight the potential therapeutic value of targeting the MT cytoskeleton and its complex regulation to combat viral infections.

## Results

### The MT-remodeler Trim69 inhibits spreading infections of lab-adapted and transmitted founder HIV-1 strains

In our previous study, we identified Trim69 as an inhibitor of the early phases of the life cycle of HIV-1 ^18^, but we had not determined its effects in spreading infections. To answer this question, we generated stable, doxycycline-inducible pools of control and Trim69-expressing TZM-BL cells. TZM-BL cells are HeLa cell derivatives expressing the CD4 receptor, the CXCR4/CCR5 co-receptors and containing an integrated LTR-Luciferase reporter for monitoring viral spread. Control and Trim69-expressing cells were treated with doxycycline to induce Trim69 expression, then challenged with replication-competent viruses. The extent of viral spread was measured over time. Under these conditions, Trim69 expression led to a profound inhibition of HIV-1 replication for both lab-adapted and transmitted founder strains (Fig. 1A), confirming that Trim69 is a *bona fide* inhibitor of HIV replicative infection. Further analysis using a single-round of infection-competent, JR-FL Env-pseudotyped HIV-1 viruses confirmed that the infectivity defect imposed by Trim69 maps to the early phases of the HIV-1 cycle (Fig. 1B). This was accompanied by an important reorganization of the MT network, similar to what we previously observed in macrophage-like THP-PMA cells (Fig. 1C) and ^18^). Specifically, Trim69 bound and bundled MTs, leading to the increase of post-translational modifications associated with stable MTs, i.e. detyrosination and acetylation of α-tubulin (Fig. 1C and Extended Data 1A for confocal microscopy and Extended Data 1B for immuno-gold EM analysis). The similar effects of Trim69 on cycling TZM-BL and growth-arrested THP-PMA cells allowed us to examine and exclude potential deleterious impacts of Trim69 on cell division, or cell survival. Indeed, under the conditions used here neither the proportion of apoptotic cells, nor the cell cycle progression, or the cell mitotic index (in the case of dividing TZM-BL cells) were affected by the presence of Trim69 during a time frame that exceeded our infection protocols (Extended Data 1C, 1D and 1E). Given their comparable Trim69-induced phenotypes, THP-PMA or TZM-BL cells were used to characterize the molecular mechanisms underlying the antiviral activity of Trim69.

**Figure 1.**
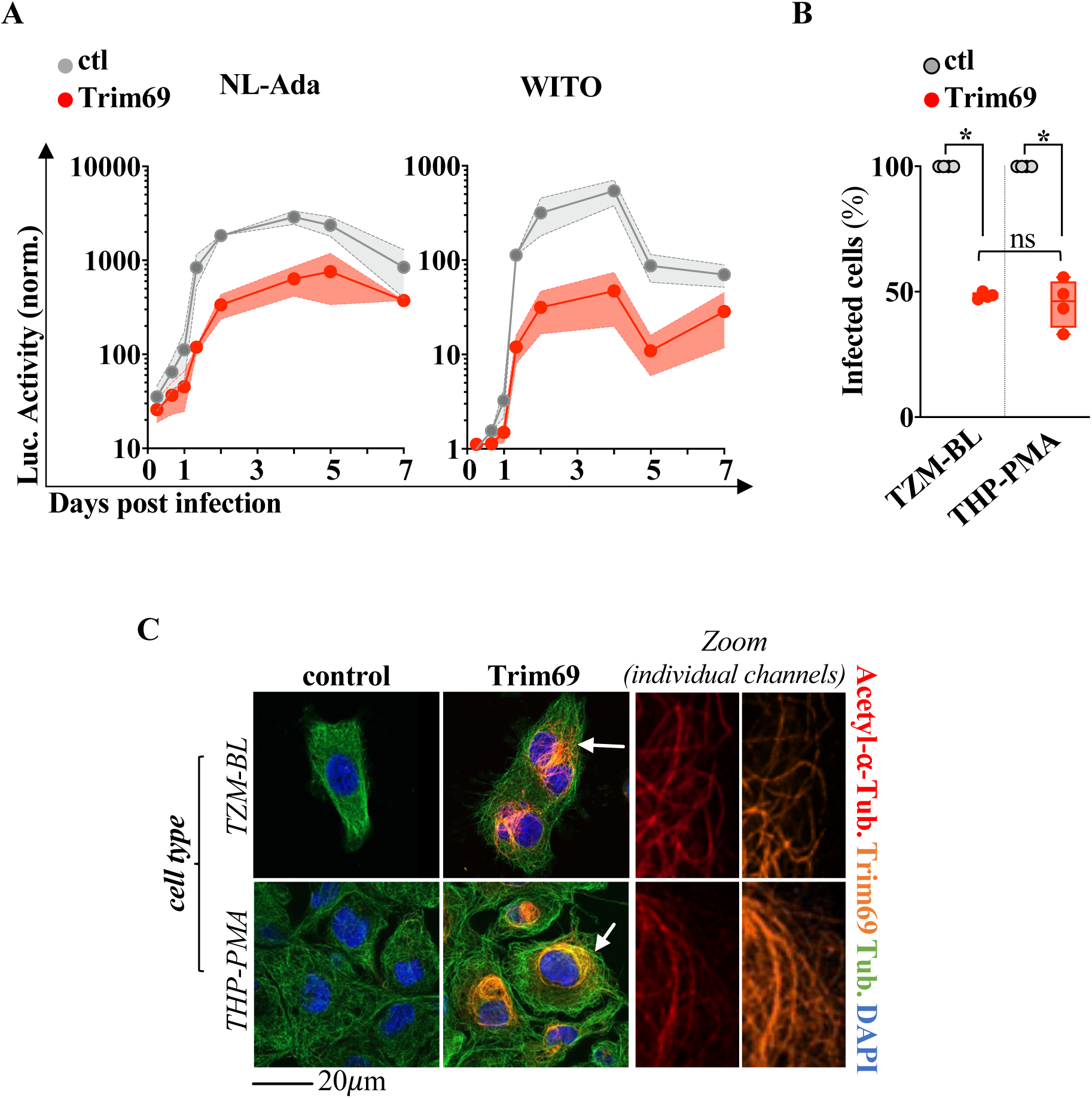
The Trim69 MT-remodeler inhibits HIV-1 viral spread. **A)** Control and Trim69-expressing doxycycline (dox)-inducible stable TZM-BL cell lines were generated via retroviral-mediated gene transduction and selection of pools of resistant cells. After dox. induction of both control and Trim69 cells, cells were challenged with the indicated HIV-1 strains (lab-adapted and transmitted founder) with multiplicities of infection (MOI) ranging from 0.01 and 0.05. Viral spread through the culture was monitored at different days post infection by quantifying Firefly Luciferase (FLuc) activity, expressed under the control of the HIV-1 LTR in TZM-BL cells. The graph presents Avg and SEM of 3 independent infections. **B and C)** as in A, but TZM-BL or stable THP-1 cells induced into a macrophage-like status by incubation with PMA (THP-PMA) were either challenged with a single-round of infection competent HIV-1 virus bearing the R5 tropic HIV-1 envelope JR-FL and coding for a GFP reporter (MOIs comprised between 0.5 and 1, B), or analyzed by confocal microscopy (C). In the case of single-round infections, the extent of infection was determined by flow cytometry two to three days post viral challenge. Avg and SEM from 4 independent experiments are shown in B, while the panels in C present typical results obtained (n>3). *, p<0.05 following an unpaired, two-tailed Student t-test between the indicated conditions.

### *In silico* modeling and TIRF-based microscopy identify the determinants of the interaction between Trim69 and MTs

To identify the specific domain/s responsible for the association between Trim69 and MTs, we used AlphaFold 2.0 structural modeling to predict the interaction of Trim69 with a single α/β-tubulin single heterodimer (Fig. 2A and Extended Data 2A, 2B and 2C ^23^, PRABI Lyon-Gerland Facility Service). ESPript analysis identified a potential interaction between a basic surface in the SPRY domain of Trim69 and the acidic C-terminal tail of the β-tubulin isotype Tubb2b (Fig. 2A and Extended Data 2A, 2B and 2C ^24^). The SPRY represents the most divergent domain among members of the Trim family and it is in general the one responsible for the binding specificity of individual Trim proteins. Interestingly, the surface of potential interaction of the SPRY domain of Trim69 and tubulin is absent in Trim5α, a well-known retroviral inhibitor that does not bind to MTs. Instead, the corresponding surface of the SPRY domain is occupied by the three variable loops that are required for interaction with retroviral capsids in Trim5α (Extended Data 2D) ^25^. Given that (1) the single α/β-tubulin dimer used in our modelling does not recapitulate the full MT structure, and (2) the C-terminal tails of all α- and β-tubulins are enriched in acidic amino acids (Extended Data 3), we investigated whether the C-terminus of α-tubulin could also bind Trim69. Structural modeling of isolated SPRY domain and the C-terminal region of α-tubulin (Tuba1a, AlphaFold webserver, ^26^) revealed that these domains could indeed potentially interact (Fig 2A). These findings suggest that the interaction between Trim69 and microtubules could be mediated by the binding of the basic pocket in the SPRY domain to the acidic C-terminal region of α- and β-tubulins.

**Figure 2.**
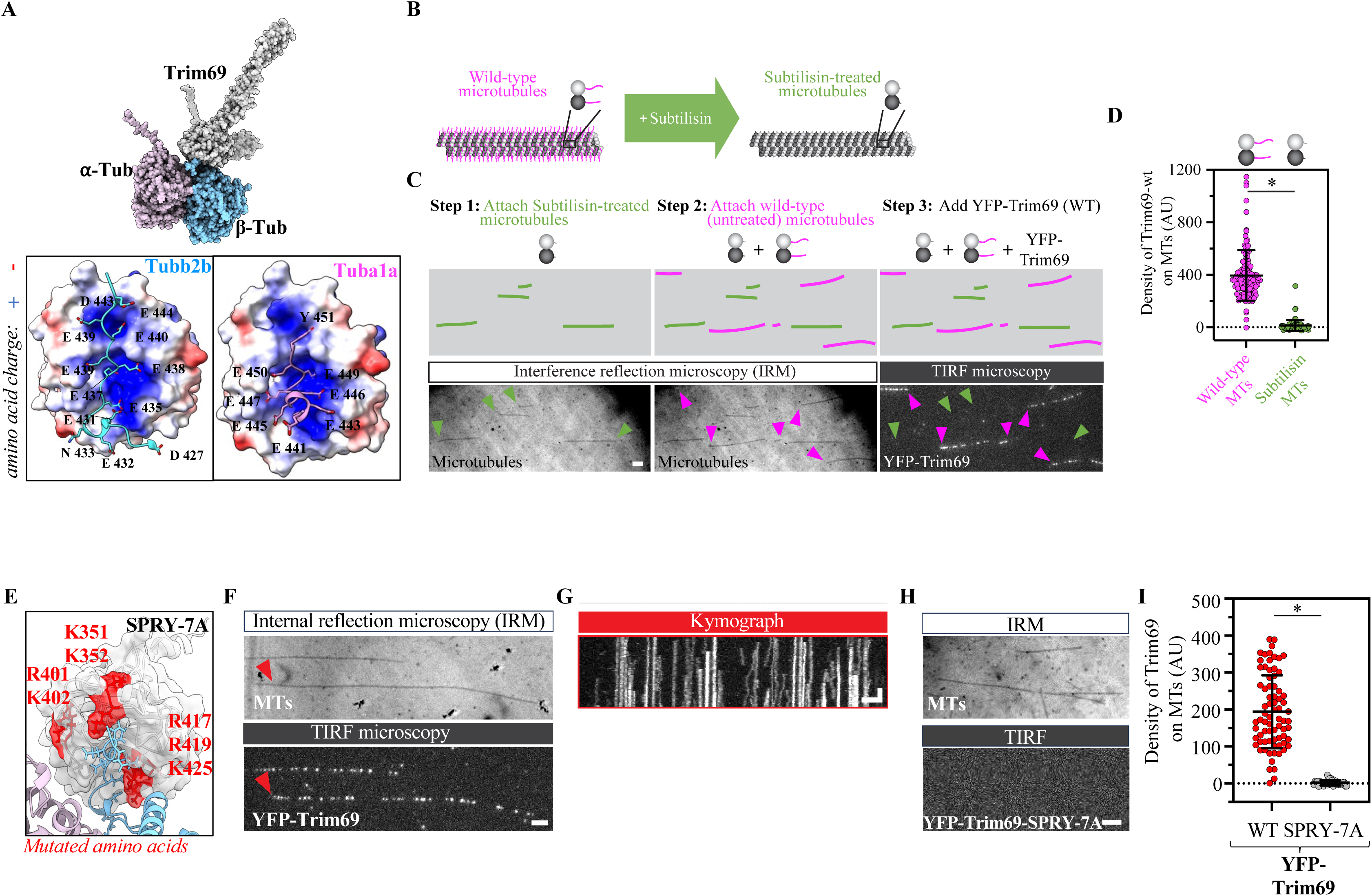
*In silico* modeling coupled to TIRF identifies the domains of interaction between Trim69 and microtubules. **A)** AlphaFold 2.0 modeling following by an ESPript analysis was used to predict complexes potentially interacting with Trim69. The inset highlights the region of interaction between the SPRY domain of Trim69 (grey), the Cter tail of β-tubulin (cyan) which was then refined with AlphaFold 3.0 also on the C-terminal tail of α-tubulin (pink). Blue/red surfaces indicate positive and negative charges respectively. **B)** Schematic representation of Subtilisin treatment of microtubules. Wild-type microtubules are incubated with Subtilisin, leading to cleavage of the C-terminal tails of α- and β-tubulin. **C)** Representation of the TIRF assay with Subtilisin-treated and untreated microtubules. IRM/TIRF images (bottom panels) and schematic drawings (top panels) identify the two microtubule types. Label-free Subtilisin-treated Taxol-stabilized microtubules (green arrowheads) and wild-type (untreated) Taxol-stabilized microtubules (magenta arrowheads) are sequentially immobilized on a glass coverslip and imaged using IRM (step 1 and 2). Next, 1000x diluted YFP-Trim69-wt-overexpressing lysate is added to the chamber and the Trim69-microtubule binding is recorded in TIRF (right panel). Scalebar: 2 µm. **D)** The dot plot presents cumulative densities of Trim69 after 3 min incubation on Wild-type (magenta) MTs or Subtilisin-treated MTs (green) in 4 independent experiments. E) Residues mutated in the Trim69 mutant SPRY-7A are specified and highlighted with a solid red surface of the SPRY domain of Trim69. F) Representative TIRF image of wild type YFP-Trim69-expressing lysate (bottom panel) diluted 1000x and incubated for 3 min on label-free Taxol-stabilized microtubules (top panel, visualized by IRM). **G)** Time-projected kymograph of the wild type Trim69 signal on the microtubule in F marked with a red arrowhead. Scalebars: vertical 2 µm, horizontal 1min. H) Representative TIRF image of mutant YFP-Trim69-SPRY-7A-expressing lysate (bottom panel) 1000x diluted and incubated as in F (top panel, MTs visualized by IRM). I) The dot plot presents cumulative densities in 6 to 8 independent experiments at 1000x lysate dilution. Scalebars: 2 µm. *, p<0.05 following an unpaired, two-tailed Student t-test between the indicated conditions.

To assess the importance of the C-terminal tails of tubulins for Trim69 binding, we used a cell lysate-based, total internal reflection fluorescence (TIRF) assay that allows the characterization of the behavior of fluorescent proteins expressed in a cell lysate at a single MT level in an *in vitro*, cell-free setting ^27,28^. We first studied the affinity of wild-type YFP-Trim69 to Subtilisin-treated MTs and compared it with the affinity to untreated MTs. Subtilisin-treatment results in MTs devoid of the protruding C-terminal tails of both α- and β-tubulins providing us with the opportunity to directly assess the importance of the tubulin tails in the binding of Trim69 (scheme of Fig. 2B, ^29^). Each type of MT was stabilized using Taxol and immobilized sequentially in the same experimental chamber where they were identified by interference reflection microscopy (IRM). Next, diluted lysates of HEK293T cells overexpressing YFP-Trim69 was added, and fluorescence was visualized by TIRF microscopy (scheme of Fig. 2C). Under these conditions, YFP-Trim69 associated to untreated MTs, but failed to bind to Subtilisin-treated MTs with cumulative fluorescent signals that dropped from 395 ± 193 (on n=87 individual microtubules) in the case of untreated MTs, to 14 ± 41 (on n=113 individual microtubules) for Subtilisin-treated ones (Fig. 2C for representative pictures, 2D for cumulative data and Expanded Movie 1A). Therefore, these results indicate that the C-terminal tails of α- and β-tubulins are required for efficient association of Trim69 to MTs.

Next, we assessed the functional role of the basic SPRY pocket in Trim69. To this end, we mutated seven basic residues that were predicted to be either in direct contact with β-tubulin (R417, R419, K425, K351), or located in flexible regions that could potentially rearrange upon MT binding (K352, R401, K402), generating the Trim69 mutant SPRY-7A (Fig. 2E). To investigate the effect of these mutations, we first compared the binding of wild-type and SPRY-7A mutant YFP-Trim69 to MTs in the cell-free, *in vitro* TIRF assay. This time a single type of MT (untreated) was immobilized in experimental chambers and equal amounts of either YFP-Trim69 or YFP-Trim69-SPRY-7A were then added to the chambers and imaged in TIRF mode (Fig. 2F, 2G, 2H for representative pictures, 2I for cumulative data and Expanded Movies 1B and 1C). Wild-type YFP-Trim69 formed distinct puncta on MTs that are mostly immobile, with few that diffused along the MT lattice (Fig. 2G). Strikingly, the SPRY-7A mutant completely failed to bind to MTs (Fig. 2H and 2I). Specifically, the YFP-Trim69 WT fluorescent intensity on microtubules was of 194 ± 99 (n=75 microtubules), while YFP-Trim69-SPRY-7A mutant showed a fluorescence of mere 2 ± 7 (n=34 microtubules). This difference was not due to a lower concentration of the SPRY-7A mutant in cell lysates, as similar results were observed over a range of different cell lysate dilutions (Extended Data 4A, 4B and 4C) and also given that this mutant was robustly expressed in cells (see below). Overall, these results demonstrate that the interaction between Trim69 and MTs is mediated by the association between a basic pocket in the SPRY domain of Trim69 and the C-terminal tails of tubulins.

### The basic pocket in the SPRY domain is the crucial determinant of MT remodeling and antiviral activities of Trim69 in cells

We next characterized the effect of mutations in the basic pocket of Trim69 in cells. When expressed in TZM-BL, the Trim69-SPRY-7A mutant exhibited a diffuse cytoplasmic localization and failed to induce MT acetylation (Fig. 3A for a WB analysis and 3B for confocal microscopy), in agreement with the observation that this mutant has lost its ability to associate with MTs. More importantly, the Trim69-SPRY-7A mutant also lost its antiviral activity and failed to protect cells from HIV-1 infection (Fig. 3C and 3D). These results therefore underscore the essential role of the basic pocket in the SPRY domain of Trim69 for MT binding, MT remodeling and antiviral activities. Of note, the SPRY-7A mutant retained its ability to dimerize as demonstrated through two complimentary approaches: first, co-immunoprecipitation with wild-type Trim69 (Extended Data 4D); second, by its ability to regain MT distribution upon co-expression with wild-type Trim69 (Extended Data 4E). These findings indicate that while the SPRY-7A mutation disrupts MT binding and the subsequent antiviral activities of Trim69, it does not impair its ability to undergo homotypic Trim69:Trim69 interactions. Finally, to further support the importance of this domain in the functions of Trim69, we performed phylogenetic analyses of the SPRY domain across mammalian Trim69 proteins (n=158 orthologous sequences). These analyses revealed that the basic character of 6 out of the 7 residues is 100% conserved in mammalian Trim69. Only the residue at position 352 was variable, being basic in all simian primates and in a couple of isolated mammalian species (i.e. *Heterocephalus glaber* among rodents and *Phascolarctos cinereus* among marsupials). These results show that the majority of the residues involved in the interaction with MTs are conserved, strengthening the hypothesis that MT-binding is a fundamental feature of Trim69 which is maintained through evolution (Extended Data 5).

**Figure 3.**
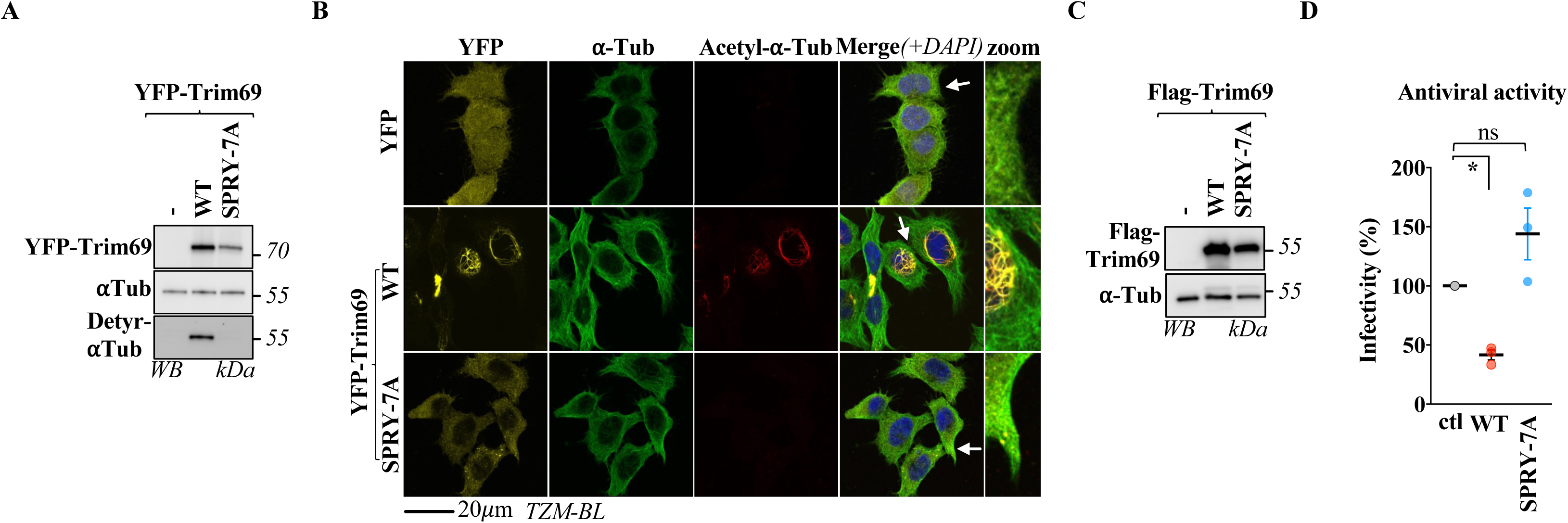
The basic surface in the SPRY domain of Trim69 is crucial for MT remodeling and antiviral activities in cells. **A and B)** WT and SPRY-7A Trim69 proteins expressed as Nter fusions to YFP were expressed in TZM-BL cells prior to WB and confocal microscopy analyses. **C and D)** Flag-tagged *wild-type* and SPRY-7A Trim69 proteins were expressed as above and cells were either analyzed by WB, or they were challenged with single round of infection-competent HIV-1 virus pseudotyped with the JR-FL envelope and coding for a GFP reporter, prior to flow cytometry analysis 2-3 days afterwards. The different panels present representative results obtained, while the graph presents Avg and SEM from independent experiments. * and ns, p<0.05 and non statistically significant, respectively as indicated following an Ordinary one-way Anova test (Sidak’s multiple comparison test).

### Trim69 decreases the number of end-binding protein 1 (EB1) comets and lowers the number of viral cores docked on MTs

EB1 comets, which specifically associate with growing plus ends of MTs, have been proposed as binding points for HIV-1 viral cores to MTs at the cell periphery ^11^. To investigate whether Trim69-mediated MT reorganization could affect HIV-1 infectivity by reducing the landing of viral cores on MTs, we quantified the number of EB1 comets in the presence and absence of Trim69 using confocal microscopy and 3D reconstruction of EB1 comets in individual cells (Fig. 4A, >1000 comets per condition across 15 to 16 individual cells per group). Similar qualitative changes in EB1 comets were observed in TZM-BL (Extended Data 6) and THP-PMA cells. However, as endogenous EB1 staining was stronger in THP-PMA cells, the full quantitative analysis was conducted in this cell type.

**Figure 4.**
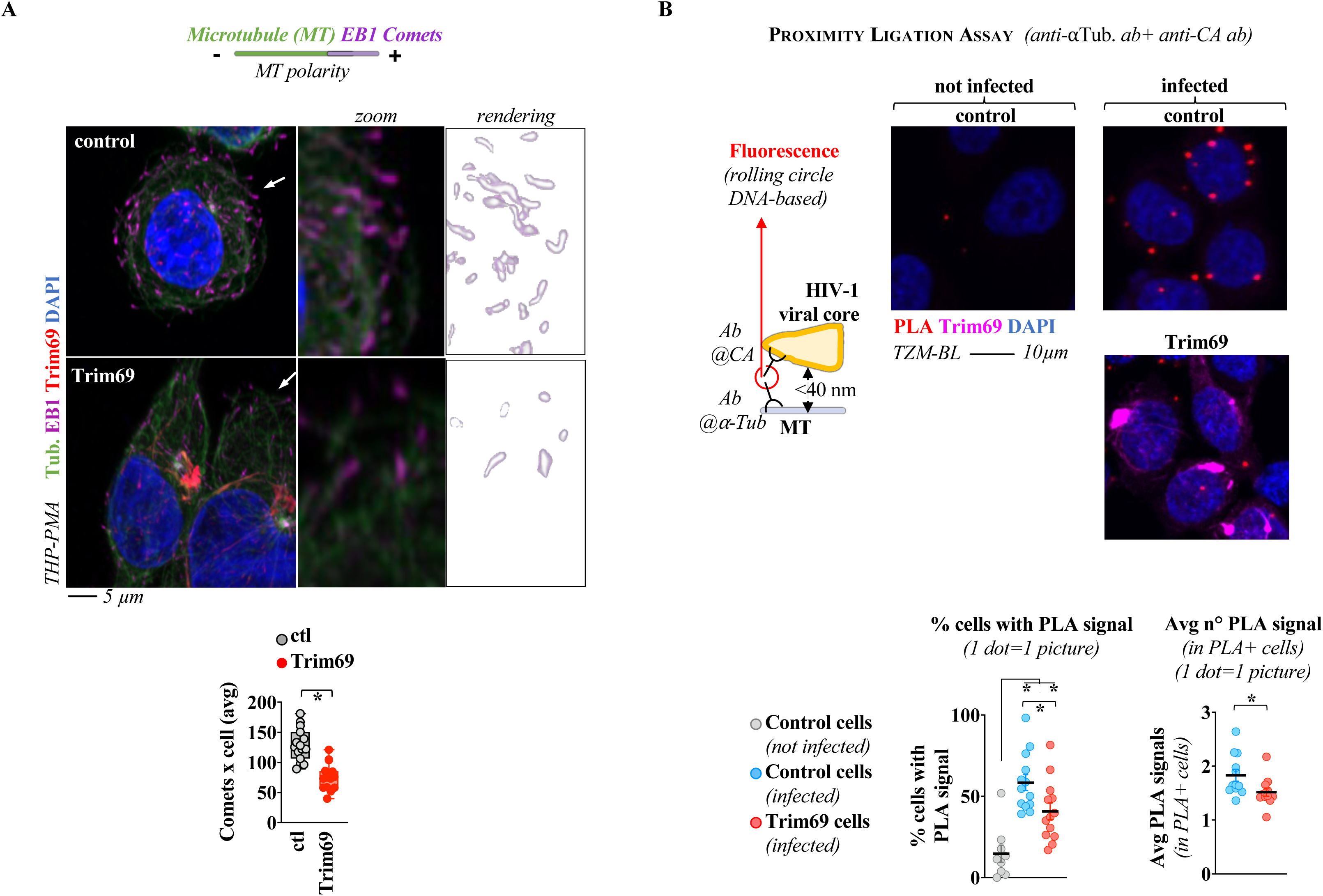
Trim69 reduces the number of EB1 comets on MT tips and decreases the docking of virion cores to MTs. **A)** THP-PMA cells expressing or not Trim69 were analyzed by confocal microscopy. After 3D reconstruction, EB1 comets were then quantified on a per cell basis. The graph presents the average number of comets per cell (>1000 comets per condition across 15 to 16 individual cells per group in 3 independent experiments). **B)** Schematic presentation of the PLA assay and antibodies used here. The panels present typical results obtained, while the graphs present the quantification of the proportion of PLA-positive cells and the quantification of the number of PLA+ dots per PLA-positive cell (> 1100 individual cells analyzed per condition in 3 independent experiments). In the Trim69 condition, only Trim69-positive cells were analyzed. Statistics were as follows: * p<0.05 following an unpaired, two-tailed, Student t-test (A and right graph in B), or an Ordinary one-way Anova, Tukey’s multiple comparison tests (left graph in B), between the indicated conditions.

Our results indicate that cells expressing Trim69 exhibit a significant decrease in the average number of EB1 comets compared to control cells, while no notable changes were observed in comet length (Fig. 4A, Extended Data 7, Expanded Movie 2). Interestingly, the distribution of EB1 in both control and Trim69-expressing cells was markedly different from the broader redistribution of EB1 along the MT shaft observed in cells treated with the microtubule-stabilizing compound Taxol. This distinction supports the notion that Trim69 and Taxol, the latter of which has been reported to exert a proviral effect in the context of HIV infection ^30^, generate phenotypically distinct subsets of stable MTs (Extended Data 7).

To determine whether Trim69 could also lead to a reduced number of HIV-1 cores docking onto MTs, we used proximity ligation assay (PLA) with antibodies specific to α-tubulin and HIV-1 Capsid (Fig. 4B), as in this assay fluorescence is generated when the two target antigens are located within at most 40 nm of each other (scheme Fig. 4B). Prior to these experiments however, we carried out infections with virions containing the Vpr-INmNeonGreen fusion protein which labels incoming virion cores ^31^ to exclude potential effects of Trim69 in virus entry, parameter that could alter the interpretation of PLA experiments. To this end, cells were challenged with virus and fixed two hours post-infection for confocal microscopy analysis. Under these conditions, >99% of cells scored positive for virion cores in both control and Trim69-expressing cells (Extended Data 8A), indicating that Trim69 does not cause a virus entry defect, confirming what we had previously reported with other techniques ^18^. PLA experiments were then carried out using the same experimental conditions and viral inputs using JR-FL-pseudotyped HIV-1 viruses (bearing no Vpr-INmNeonGreen). The number of viral cores docked on MTs (PLA-positive dots) was quantified on a per cell basis (> 1100 individual cells analyzed per condition; in the case of Trim69, the analysis was carried out only in Trim69-positive cells). Under these conditions, both the number of cells bearing PLA-positive signal, as well as the number of PLA dots per positive cell decreased significantly in presence of Trim69 (averages of 58.4% vs 40.7% and 1.83 vs 1.51, respectively, Fig. 4B and Extended Data 8B). Given that this decrease is not due to an entry defect, these results indicate that Trim69 leads to a decrease in the number of virion cores that successfully dock on MTs.

### Trim69 functionally mirrors dynein motor inhibition during HIV-1 infection

To determine whether Trim69 inhibits HIV-1 infection by interfering with the dynein-dependent migration of viral cores, we compared the effects of Trim69 overexpression with dynein inhibition during HIV-1 infection. Dynein inhibition was achieved either chemically, using Ciliobrevin D (Extended Data 9A), or genetically, through the silencing of two key dynein complex components: the dynein heavy chain 1 (DYNC1H1) and the bicaudal D cargo adaptor 2 (BICD2, Fig. 5A). Under both conditions, HIV-1 infection was impaired in control cells (Extended Data 9A, Fig. 5A, respectively and Extended Data 9B for silencing efficiencies) and interestingly, the extent of HIV-1 inhibition caused by dynein inhibition was similar and non-additive to the one observed by expression of Trim69, indicating that Trim69 and dynein operate along the same pathway.

**Figure 5.**
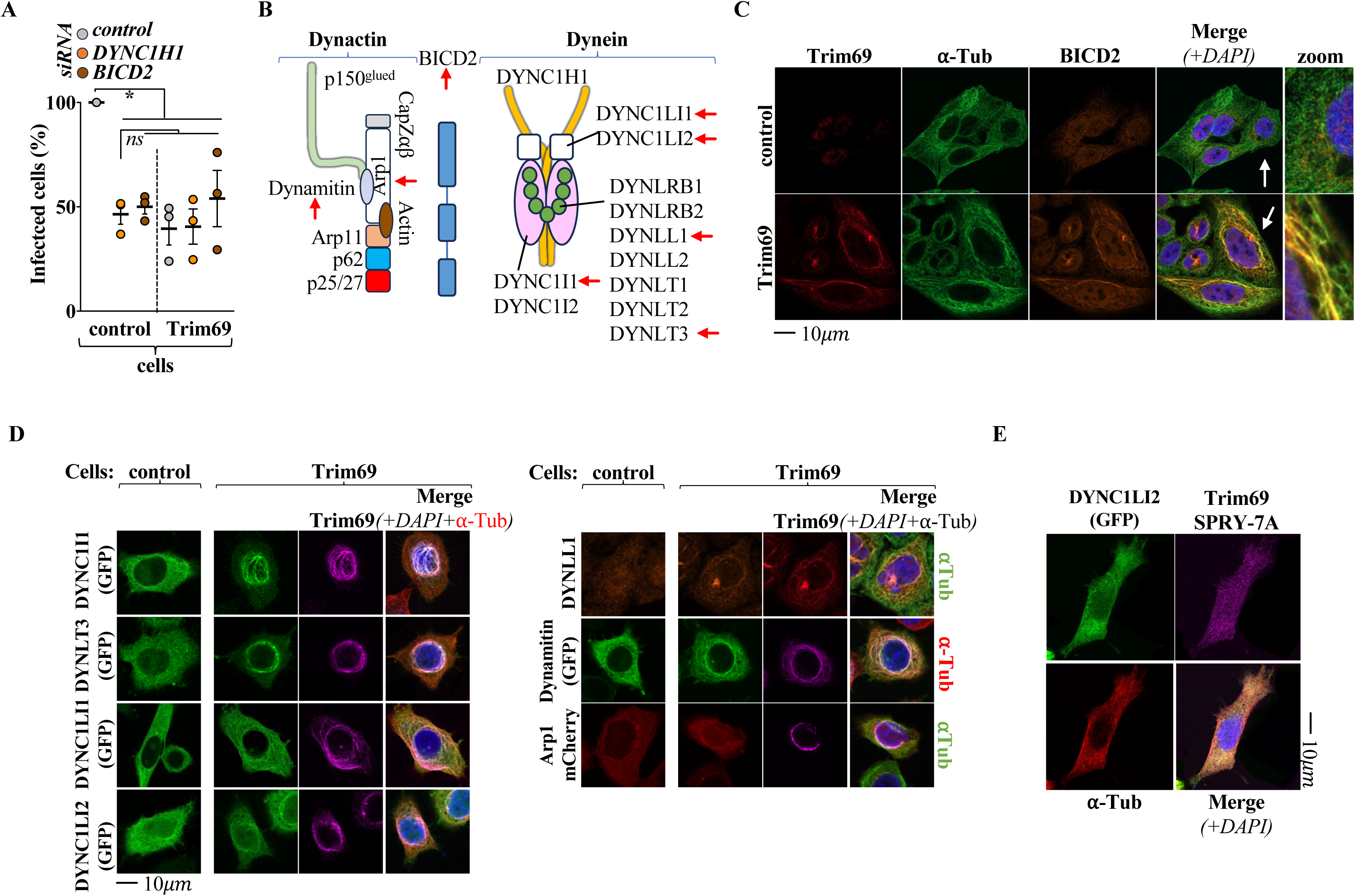
Trim69-mediated inhibition of HIV-1 infection mimics the effects of dynein inhibition and Trim69 relocalizes dynein and dynactin complexes to Trim69-decorated MTs. **A)** TZM-BL cells expressing or not Trim69 were challenged with GFP-coding and JR-FL Env-bearing single round of infection-competent HIV-1 virus following siRNA transfection of the indicated proteins (efficacy of silencing presented in Extended Data 9B). The extent of infection was measured by flow cytometry two to three days later. **B)** Schematic organization of the dynein and dynactin complexes along with the BICD2 adaptor. Red arrows indicate the components analyzed by confocal microscopy using either antibodies directed against endogenous proteins (**C)** BICD2 and Extended Data 10 for DYNLL1), or using ectopic transfection of GFP- or mCherry-fusion components (**D**). The complete panel analysis is provided in the Extended Data 10). **E)** TZM-BL cells expressing or not DYNC1LI2 along with the SPRY-7A Trim69 mutant were obtained by transient transfection. Cells were then analyzed by confocal microscopy. The graph present averages and SEM while panels present typical images obtained (n= 3). * and ns, p<0.05 and non-significant following one-way Anova, Tukey’s multiple comparison tests between the indicated conditions.

### Trim69 causes stalling of the dynein and dynactin motor complexes on MTs

Cargo transport towards the MT minus-end relies on the proper assembly of dynein and dynactin complexes with specific adaptors ^32^ (schematically illustrated in Fig. 5B). To determine whether Trim69 affects these complexes, we examined the spatial distribution of eight dynein and dynactin components, used as markers for the whole complexes, by confocal microscopy. These components included endogenous proteins such as BICD2 (Fig. 5C) and DYNLL1 (Extended Data 10) detected via antibody staining, as well as ectopically expressed GFP- or mCherry-tagged fusion proteins (Fig. 5D and Extended Data 10). In all cases, components that were dispersed in control cells became redistributed on Trim69-decorated MTs (Fig. 5C and 5D, and Extended Data 10 for individual channels). This redistribution was specific to Trim69 and not a general consequence of MT stabilization, as no accumulation was observed in cells treated with MT-stabilizing agent Taxol (Extended Data 11). To confirm that the accumulation of dynein/dynactin components was functionally important in the antiviral effect of Trim69, we evaluated the behavior of the non-functional SPRY-7A Trim69 mutant with DYNC1LI2 (Fig. 5E). In contrast to *wild-type*, the SPRY-7A Trim69 mutant failed to induce an accumulation of DYNC1LI2, in line with the loss of function of this mutant.

Altogether, these results demonstrate that Trim69 induces the redistribution and accumulation of dynein and dynactin complexes along microtubules to which it is bound.

### Trim69 is a strong MTs binder

We hypothesized that the accumulation of motor complex components on MTs decorated with Trim69 could be explained by a strong association of Trim69 with MTs that physically impedes the passage of dynein:cargo complexes. To determine whether this was the case, we performed fluorescence recovery after photobleaching (FRAP) experiments both *in vitro* on isolated MTs using TIRF microscopy, as well as in cells. In this setting, Trim69 molecules are subjected to photobleaching, causing them to lose fluorescence. The time of fluorescence recovery informs about the dynamic properties of the protein and the strength of the interaction between the protein and the MT. If a protein binds dynamically or weakly, the molecules which lost the fluorescence will be rapidly replaced by new, still fluorescent ones. Conversely, proteins that associate irreversibly, or strongly to the MT will remain bound and no fluorescence recovery will be observed. In the *in vitro* TIRF experiment, wild-type YFP-Trim69 was incubated with Taxol-stabilized MTs and allowed to form puncta on MTs, as observed before (Fig.2F). We then photobleached a rectangular region of the Trim69-covered MT and studied the recovery of the Trim69 signal along the MT over time. The resulting recovery curve revealed a poor fluorescent recovery of the Trim69 signal within the experimental timeframe (Fig. 6A, 6B and 6C for representative pictures, corresponding kymograph and a graph for cumulative data and Expanded Movie 3A). Specifically, we found a 20 ± 23% fluorescence recovery at 6.5 min after photobleaching, however, this recovery could largely be attributed to new Trim69 binding events to the MT lattice (marked with red stars in Fig. 6B), rather than to the replenishment of pre-existing Trim69 puncta, indicating a low turnover and strong association of Trim69 with MTs. Similarly, FRAP experiments were conducted in cells expressing mScarlet-Trim69 where a random MT-rich area within the cell was subjected to photobleaching and the fluorescence recovery was evaluated by live imaging (Fig. 6D for representative pictures, 6E for cumulative data and Expanded Movie 3B). These experiments showed a lack of restoration of Trim69 fluorescence after photobleaching, indicating again that Trim69 is strongly associated to MTs in cells. Combined, this data supports our hypothesis that Trim69 may act as a roadblock on MTs that leads to the accumulation of dynein motor complexes.

**Figure 6.**
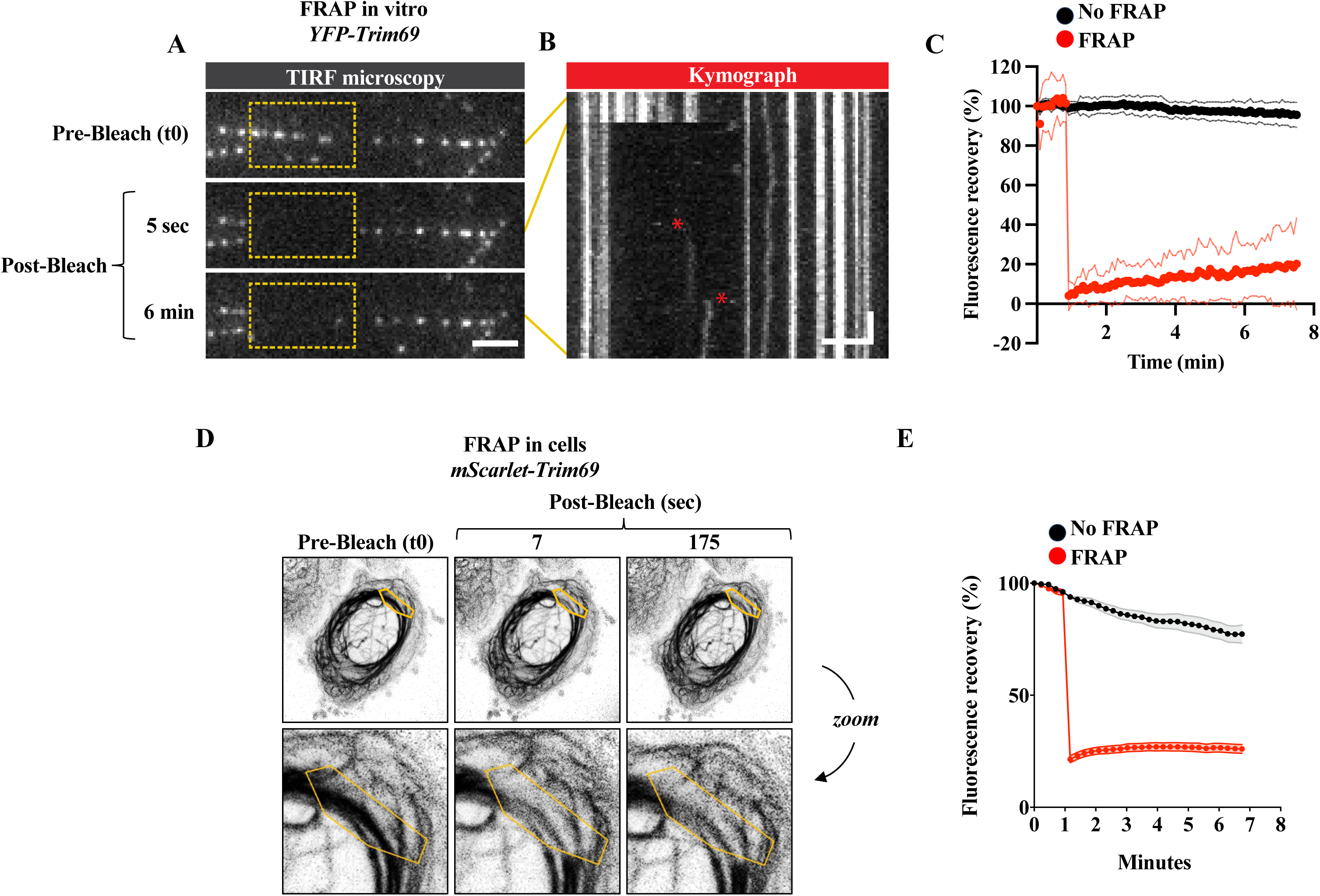
FRAP experiments indicate a strong association between Trim69 and MTs both *ex vivo* and in cells. **A)** Representative TIRF images of wild-type YFP-Trim69 expressing lysate, 1000x diluted and incubated on Taxol-stabilized MTs. Rectangular regions along the MT lattice were subjected to photobleaching (area marked with yellow box), after which the recovery of the fluorescent signal was followed for 6.5 min. Scalebar: 2 µm. **B)** Time-projected kymograph of the MT shown in A. New Trim69 binding events are marked with red stars. Scalebars: vertical 2 µm, horizontal 1 min. **C)** Cumulative graph showing the fluorescence recovery of Trim69-wt signal on 24 MTs (FRAP, red) and 22 non-FRAP MTs (non-FRAP, black). **D and E)** Similar experiments were conducted in TZM-BL cells expressing an mScarlet-Trim69 protein following live imaging (representative images and cumulative graphs from 20 individual cells analyzed, respectively).

### Trim69 induces an overall migration defect of viral cores towards the nucleus

To determine whether Trim69 could impair the dynein-dependent migration of virion cores towards the nucleus, we first decided to examine the migration behavior of mitochondria, which are well-known dynein-dependent cargos ^33^. Live imaging and trajectory reconstruction of individual PKmito-labeled mitochondria showed a marked reduction in the distance travelled and mean speed in cells expressing Trim69 (Extended Data 12A and 12B, data from 7-10 individual cells and between 4700-7500 individual trajectories per condition), strongly supporting the possibility that Trim69 acts as a physical barrier for the dynein motor on MTs. To determine whether this was the case also for HIV-1 virion cores, we tracked the migration of single virion cores along MTs by live imaging. Cells were infected with VSVg -pseudotyped, Vpr-INmNeonGreen protein-containing virus ^31^ for 2 hours and MTs were visualized using SiR-Tubulin (Fig. 7A for representative pictures and graph for analysis of >250 individual cores analyzed across eight cells per condition; Extended Data 12C for a lower magnification view of the cells and Expanded Movie 4). Under these conditions, we determined that Trim69 inhibits the dynamic movement of virion cores along MTs with track displacement rates that passed from 0.970 μm in control cells to 0.425 μm in the presence of Trim69.

**Figure 7.**
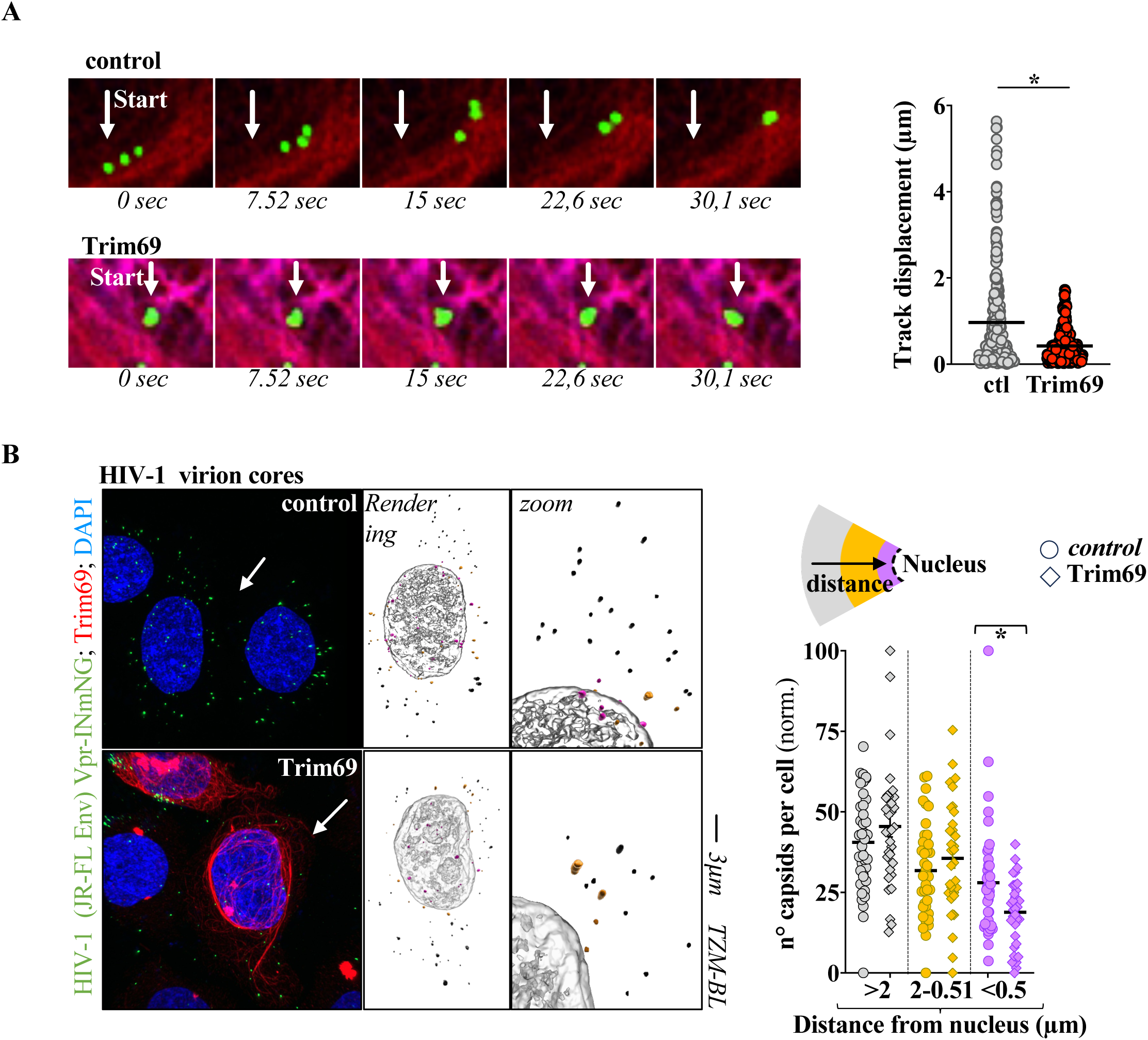
Trim69 impairs dynein-dependent migration of HIV-1 virion cores towards the nucleus. TZM-BL cells expressing or not Trim69 were challenged with an HIV-1 virus pseudotyped with the VSVg envelope and incorporating a Vpr-Neongreen-IN that labels virion cores. **A**) MTs were labeled for 1 hour upon incubation with 1µM of SiR-Tubulin with 10 µM Verapamil and the movement of virion cores was tracked by live imaging. The pictures present representative results obtained (lower magnification images are provided in the Extended Data Fig. 12A), while the graph presents the track displacement analysis of >250 individual cores across eight cells per condition. **B**) Cells were analyzed two hours post infection cells prior to confocal microscopy and 3D reconstruction. The pictures present representative distributions. The graph presents Avg and SEM obtained from the analysis of over 3300 viral cores in more than 30 cells across four independent experiments. Viral cores are color-coded according to their distance to the nucleus in both representative rendered pictures and graph. *, p<0.05 following an unpaired, two-tailed Student t test between the control and Trim69 indicated conditions.

Given that the measured total distances travelled in the presence of Trim69 similarly dropped from 2.284 μm in control cells to 1.359 μm in cells expressing Trim69, we next decided to assess whether this could translate into an overall defect of migration of HIV-1 virion core populations as they approach the nucleus. We thus analyzed the spatial distribution of fluorescently-labelled HIV-1 virions in TZM-BL cells with or without Trim69 expression with respect to the nucleus. To this end, cells were infected with Vpr-INmNeonGreen protein-containing virus for two hours, fixed, and the position of viral cores relative to the nucleus were quantified using confocal microscopy and 3D cell reconstruction. While no differences were observed at distal locations, cells expressing Trim69 exhibited a significant reduction in the number of cores near the nucleus from 14.01% to 9.43% of the total number of virion cores present in the entire volume of the cell (Fig. 7B for representative images and quantitative analysis from over 3300 viral cores in more than 30 cells across four experiments and Expanded Movie 5). Thus, Trim69 impairs retrograde transport of HIV-1 virion cores along microtubules, inhibiting their overall progression and accumulation in the proximity of the nucleus.

Taken together, these results demonstrate that Trim69-mediated MT remodeling inhibits dynein-dependent cargo transport of HIV-1 virion cores along MTs towards the nucleus.

## Discussion

This study demonstrates that Trim69 impairs HIV infection by disrupting the dynein-dependent transport of virion cores through a profound remodeling of the MT network. This reorganization leads to a reduction in the number of dynamic MTs, as evidenced by fewer EB1 comets, to a decrease in the docking of virion cores on MTs and to the inhibition of dynein-dependent cargo movement. These effects concur in decreasing the proportion of virion cores that reach the nuclear periphery, effectively inhibiting HIV-1 infection. The reorganization of MTs by Trim69 relies on a positively charged pocket within its SPRY domain, which is essential for binding to MTs through its interaction with the C-terminal tails of tubulins and is critical for the antiviral function of Trim69.

The MT network is a major axis of transportation within the cell and it is thus not surprisingly used by many viruses to accomplish their life cycle. Several MT-interacting proteins (MAPs, motors, adaptor proteins) have so far been identified as facilitators of viral infection ^2-13^. However, far less is known about whether the MT cytoskeleton can be actively reprogrammed to inhibit viral infection. To our knowledge, Trim69 is the first example of innate immune factor capable of inducing a profound reorganization of the MTs and to inhibit viral infection ^18^, thus highlighting the impact that MT remodeling may have in viral control.

Trim69-induced MT remodeling shares some classical features of MTs stabilized by Taxol, such as resistance to nocodazole and accumulation of high levels of acetylation and detyrosination of α-tubulin^18^. However, they also show several notable differentiating features that likely relate to the manner in which Trim69 binds and reorganizes MTs. Unlike the Taxol treatment, Trim69 overexpression in cells causes unique patterns of distribution of EB1 comets, distinct localization of dynein and dynactin complexes, and an opposite impact on viral infection, because while Taxol enhances HIV-1 infection ^2,18,30^, Trim69 inhibits it. A recent study showed that acetylation is not important for the antiviral effects of Trim69 ^22^, adding to the long-discussed question as to whether post-translational modifications of α-tubulin accumulate as cause or consequence of microtubule stabilization.

Trim69 can drive MT reorganization through multiple and not mutually exclusive mechanisms: by modulating the activity of enzymes involved in the regulation of dynamic and stable MT pools ^17^; by directly binding to, bundling and thus stabilizing MTs; or a combination of both. Our FRAP experiments indicate that the association of Trim69 with MTs is particularly stable, suggesting that Trim69 may directly stabilize MTs by multimerizing on their surface. In support of this hypothesis, MTs bearing increased levels of acetylation and detyrosination of α-tubulin are consistently decorated with Trim69. Although the extent to which Trim69 forms higher order multimeric structures remains unclear, the observation that the SPRY7A mutant, which lacks MT-binding capacity on its own, can regain a MT distribution by associating with wild-type Trim69, indicates that Trim69 multimerization is an important parameter for MT binding and reorganization.

Our results point to a model of inhibition where Trim69 creates a physical barrier on MTs that impedes the passage of dynein-cargo complexes and leads to a broad inhibition of retrograde cargo transport. If this model is correct, Trim69 should similarly interfere with kinesin-mediated transport and anterograde cargo movement, as the results obtained with mitochondria suggest. Although this hypothesis was not specifically explored in this study, such interference could further enhance the antiviral effects of Trim69 against HIV-1, given that certain kinesins have been previously involved in facilitating HIV-1 infection ^7,9,13^. Trim proteins are often endowed with E3-ubiquitin ligase activity through their RING domain. In the case of Trim69, the precise weight that this activity may have in the functions of Trim69 remains controversial ^19,21^, due to the difficulty in engineering mutants that alter the E3-ligase functions of the RING domain without affecting its overall dimeric module structure ^34^. While this topic has not been studied here, we believe it is possible that a prolonged pausing in the vicinity of Trim69 may lead to an increased ubiquitination of dynein/dynactin complex components which may constitute an additional level of Trim69-dependent regulation of motor-complexes behavior.

Trim69 interacts with MTs through a basic pocket in its SPRY domain that is highly conserved among mammalian Trim69 proteins. However, Trim69 is not the only members of the Trim family that interacts with microtubules. Several other Trim proteins and specifically MID1-Trim18, MID2-Trim1, Trim9, Trim36, Trim46 and Trim67 have also been identified as MAPs ^35-40^. For these proteins the association with MTs occurs via a loosely conserved domain termed the C-terminal subgroup one signature (COS), which is absent in Trim69 ^40^ and reviewed in ^41^. While MID1/2 have been genetically linked to a rare congenital disorder resulting from embryonic midline structures developmental defects (Opitz syndrome) ^38,42^, the pattern of expression of these Trims in immune cells, and their potential antiviral functions remain essentially unknown. The presence of multiple MAPs within the Trim family raises the possibility that the regulation of MTs by these proteins could play a more intricate role in immune responses, potentially remodeling the MT cytoskeleton towards an antiviral state. This could represent a novel layer of complexity in antiviral responses that remains to be explored and that could offer novel strategies of broad antiviral intervention.

## Methods

### Cell culture and DNA constructs

Human monocytic THP-1 (ATCC cat. TIB-202), human kidney epithelial HEK293T (ATCC cat. CRL-3216) and TZM-BL cells (obtained through the NIH HIV Reagent Program, Division of AIDS, NIAID, NIH and contributed by Dr. John C. Kappes, Dr. Xiaoyun Wu and Tranzyme Inc. cat. ARP-8129) were maintained in complete Dulbecco’s Modified Eagle Medium (DMEM, Gibco, cat. 61965-026) supplemented respectively with 10% fetal bovine serum (SERANA cat. SFBSSA015) for THP-1 and 7,5% for HEK293T and TZM-BL and 100U/mL of penicillin-streptomycin (Gibco, cat. 11548876). TZM-BL cells are a HeLa cell derivative that expresses the HIV-1 receptor/co-receptors CD4 and CXCR4/CCR5 in addition to integrated b-galactosidase and Luciferase genes under the control of an HIV-1 promoter ^43^. THP-1 cells media was also supplemented with 0.05 mM β-mercaptoethanol (ThermoFisher Scientific, cat. 31350-010), 10 mM HEPES (ThermoFisher Scientific, cat. 15630056), minimum essential medium (ThermoFisher Scientific, cat. 11140-035) and 1mM sodium pyruvate (ThermoFisher Scientific, cat. 11360-039) and macrophage-like differentiation was induced by treatment with 100 ng/ml of phorbol 12-myristate 13-acetate (PMA) (Sigma cat. P1585) for twenty-four hours.

Different N-terminal tagged versions of wild-type Trim69 were used in this study (Flag as in ^18^; GFP, YFP or mScarlet, this study) and the Trim69 SPRY-7A mutant was obtained by gene synthesis (Genewiz). Experiments were carried out using either transient ectopic DNA transfection (HEK293T cells and in some experiments with TZM-BL cells) using an in-house Calcium-Phosphate (Calcium-HBS) method, or JetPrime Transfection reagent (Polyplus, cat. POL101000015) for confocal microscopy. Alternatively, stable cells were generated by retroviral-mediated gene transduction (pRetroX-tight-Puro-based vector system (Clontech, cat. PT3960-5, see below). In this case, the expression of Trim69 was under the control of doxycycline (Sigma, cat. D5207) and dox. was of course added equally to control- and Trim69-expressing cells. Plasmids expressing the following dynein and dynactin complex components were also obtained through Addgene: DYNC1LI1 and DYNC1LI2 respectively pLVXPuro-mGFP-LIC1/-LIC2 (cat #66600 and #66601). DYNC1I1 (pCB168 cat #37389), Dynactin DCTN2 (pCB163 cat #37388), DYNLT3 pCB169 (cat #37390) in the backbone pBABE and tagged EGFP and ARP1 (PTK5 in pEGFP-C1 tagged mcherry, cat #37391).

The HIV-1 proviral constructs coding for the lab-adapted NL4-3 and the WITO transmitted founder virus were obtained through BEI Resources, NIAID, NIH: ARP-114 and HRP-11739, respectively.

### Antibodies

The following primary antibodies were used for WB, or confocal microscopy: mouse monoclonal antibodies; anti-α-tubulin, anti-Flag (Sigma cat. T5168, cat. F3165, respectively); rabbit antibodies : polyclonal anti-tubulin (Sigma, cat. SAB4500087), polyclonal anti-BICD2 antibody (Abcam, cat. ab117818), recombinant monoclonal DYNLL1/PIN antibody (Abcam, cat. ab51603); anti-detyrosinated alpha-tubulin (AdipoGen cat. REV-31-1335-00-R100) and anti-acetyl (K40)-α-tubulin (Abcam cat. ab179484) that were used interchangeably to label modified microtubules, anti-GFP (Sigma cat. G1544) and anti-phospho-histone H3 (Ser10) (Cell Signaling cat. 53348T), goat polyclonal anti-Flag (Novus Biologicals cat. NB600-344) and goat polyclonal anti-Trim69 (C-terminal) (Sigma cat. SAB2501878) antibodies, and rat monoclonal anti-MAPRE1/EB1 (Abcam ab53358) antibody. For PLA, primary antibodies anti-p24 mouse AG3.0 and 183-H12-5 were obtained through the NIH HIV Reagent Program, Division of AIDS, NIAID, NIH and were contributed by Dr. Marie-Claire Gauduin and Dr. Bruce Chesebro and Kathy Wehrly, respectively (ARP-4121 and ARP-3537, used at a concentration of 1/200).

The following secondary antibodies were used for WB: anti-mouse and anti-rabbit IgG-Peroxidase conjugated (Sigma cat. A9044 and cat. AP188P, respectively), polyclonal rabbit anti-goat Immunoglobulins HRP (Dako cat. P0449). For confocal microscopy, the following secondary antibodies were used: donkey anti-mouse IgG–Alexa Fluor 488 conjugate (Life Technologies cat. A-21202), goat anti-rabbit IgG-Alexa Fluor™ Plus 488 (Invitrogen cat. A-32731) or goat anti-rat FITC (Biorad cat. STAR69) accordingly; donkey anti-mouse or anti-rabbit IgG–Alexa Fluor 555 conjugate (Life Technologies cat. A31572 cat. A32794), or donkey anti-goat IgG–Alexa Fluor 546 conjugate (Life Technologies cat. A-11035), donkey anti-rabbit IgG–Alexa Fluor 594 conjugate and (Life Technologies cat. A-21207), donkey anti-rabbit IgG–Alexa Fluor 647 conjugate (Life Technologies cat. A-21447), or goat anti-mouse or-rabbit Alexa fluor 680 (Invitrogen cat. A-21057 and cat. A-21076). Dapi (ThermoFisher Scientific cat. 62248) was used to stain the nuclei.

### Replicative and single-round of infection competent HIV-1 virus production and infections

Replication-competent HIV-1 virus was produced by calcium phosphate transfection of HEK293T cells with DNAs coding the entire viral coding sequence in the form of a proviral construct (10 µg for 10 cm plate dish). The day after transfection, cells were washed and further cultured for twenty-four hours prior to virus harvest. The virus was quantified by challenging TZM-BL with virus dilutions and by quantifying blue infected cells using Beta galactosidase assay.

These viruses were used for viral spreading assays on TZM-BL cells. In this case, multiplicities of infection (MOIs) comprised between 0,01 and 0,05 were used to challenge cells stably expressing Trim69 or not and the extent of infection was measured at different days post infection by Luciferase assay (Promega cat. 1501).

Single-round of infection-competent HIV-1 particles were obtained similarly by transfecting four plasmids into HEK293T cells: Gag-Pro-Pol 8.2 (8,5 μg), HIV-1 reporter genome (8,5 μg); JR-FL Env (2 μg) and Rev (1 μg) for a total of 20 μg per 10cm plate using calcium phosphate method. Virus was harvested 36-48 hours after transfection, syringe-filtered (0.45 µm), ultracentrifuged at 28000 rpm at 4°C for 1h30 through a 25% sucrose cushion, resuspended, aliquoted and frozen. In this case the infectious titer was measured by infecting target TZM-BL cells with virus dilutions and by determining the percentage of GFP-positive cells two to three days post infection by flow cytometry. MOIs comprised between 0.5 and 1 were then used for the different experiments as indicated. Viruses used in proximity ligation assays (PLA) were produced similarly except that their viral genome coded for a non-fluorescent reporter (CMV:CD8), to avoid interference with the imaging procedure. When indicated, the cytoplasmic Dynein inhibitor, Ciliobrevin D (Sigma cat. 250401) was used at 20μM, one hour prior and then during infection. Viruses used in virion core detection were produced as described above, except that JetPrime was used as transfection reagent to increase transfection efficiency. Slightly modified plasmid ratios were used together with the addition of a plasmid coding for a Vpr-INmNeonGreen (Vpr-INmNG) fusion protein. Ratios were: Gag-Pro-Pol 8.91 (10 μg), HIV-1 reporter genome with CD8 gene (3,5 μg); JR-FL Env (2 μg) and Rev (1 μg) and Vpr-INmNG (4μg) per 10cm plate. The Vpr-INmNG fusion protein is incorporated into virion cores (^31^, gift from Ashwanth C. Francis, Florida State University, USA). After ultracentrifugation, virus was resuspended in FluoroBrite DMEM (without phenol red, Gibco cat. A1896701). Control or Trim69-expressing stable TZM-BL cells were seeded at 60000 cell/well in µ-Slide 8 Well high Glass Bottom (IBIDI, cat. 3507901) and treated with 1.5 μg/mL of doxycycline for eighteen hours to induce the expression of trim69. Medium was changed to FluoroBrite and cells were then infected for two hours in the presence of the reverse transcription inhibitor Nevirapine (at 10 μg/mL, cat. HRP-4666, obtained through the NIH HIV Reagent Program, NIAID, NIH).

### Generation of dox-inducible stable cell lines

Murine leukemia virus (MLV) retroviral particles were obtained by ectopic DNA calcium phosphate transfection of HEK293T cells with plasmid DNAs coding Gag-Prol-Pol (pTG5349 ^44^; the envelope protein G from the Vesicular Stomatitis Virus (VSVg); and the mini-viral genome coded by the pRetroX-Tight-Puro expressing either Trim69 or not (8:4:8 µg DNA per 10 cm plate dish, respectively). A second virus was produced using the pRetroX-Tet-On (Clontech cat. 632104) plasmid that codes for the TetOn protein and allows for doxycycline (dox)-inducible expression. Viral particles released in the supernatant of transfected HEK293T cells were harvested twenty-four to forty-eight hours after transfection and syringe-filtered (0.45 µm). Target cells were first transduced with two types of particles (pRetroX-Tight-Puro/gene of interest and pRetroX-Tet-On), and were then selected as a pool upon incubation with Puromycin (1,5 µg/ml) and G418 (1mg/ml) (Sigma cat. P8823 and ThermoFisher Scientific cat. 10131-027). Trim69 expression was routinely induced upon incubation of cells for 18 to 48 hours with 1.5 µg/mL of doxycycline (Sigma cat. D5207).

### Confocal microscopy

Cells were plated on coverslips coated with 0.01% poly-L-lysine (Sigma, cat. P4832) prior to transfection (JetPrime Polyplus cat. POL101000015), or dox induction. Cells were analyzed twenty-four hours after either transfection or induction. For analysis, cells were washed thrice with PBS 1x, fixed with 4% paraformaldehyde (15 min, Euromedex cat. 15713), quenched with 50 mM NH4Cl (10 min, Sigma cat. A4514) and permeabilized with PBS–0.5% Triton X-100 (5 min, Sigma, cat. X100). After a blocking step in PBS–3% BSA, cells were incubated with primary antibodies for 1 hour at room temperature (dilution 1:200 to 1:500), washed and then incubated with fluorescent secondary antibodies (dilution 1:200 to 1:1000) prior to incubation with 1:10000 DAPI for 10 min (ThermoFisher Scientific cat. 62248) and final mounting with Fluoromount-G™ (ThermoFisher Scientific cat. 00-4958-02). Images were acquired using a spectral Zeiss LSM800 confocal microscope or Confocal Zeiss LSM980-AiryScan and analyzed with Fiji software (version 2.0.0).

For EB1 comet analysis, THP1-PMA-differentiated cells were fixed in cold MetOH for 10 min before permeabilization ^18^. Anti-EB1 rat, Anti-Flag rabbit and anti-tubulin mouse antibodies were used at 1:200. Z-stacks were performed on Confocal Zeiss LSM980-AiryScan with the “optimal” slices option. On Imaris 9.2, surfaces were created for comets following the parameters of the algorithm available in the supplementary data. The length of each comet, as well as the number of comets per cell, was assessed with BoundingBoxOO Length C. When indicated, Taxol was added at 20uM for 24hrs.

### Virion cores detection and live imaging

For HIV-1 virion cores detection, cells were challenged with JR-FL-Env pseudotyped, Vpr-nG-IN virus for two hours. Cells were then fixed with a protocol preserving the cytoskeleton ^27^. consisting of a 10min incubation in warm Hank’s Balanced Salt Solution (HBSS), 10min in microtubules stabilizing buffer containing 1mM DSP (3,3ʹ-Dithiodipropionic acid di(N-hydroxysuccinimide ester, Sigma, cat. D3669, previously diluted in DMSO at 50mM), 15min in 4% PFA microtubule stabilizing buffer followed by a permeabilization step in 0,5% triton X-100 in stabilizing buffer. The MT stabilization buffer is composed of 1mM EGTA, 4% PEG 8000, 100mM PIPES pH 6,9. The subsequent saturation and antibodies labelling were conducted as described above in the confocal microscopy section. Imaging was performed in PBS on a confocal Zeiss LSM980 microscopy and images were analyzed on 9.8 IMARIS software. Surfaces for the viruses and the nucleus were created and the shortest distance between the surface of each particle and the nucleus was measured.

PLA staining was carried out following the supplier’s protocol (Naveniflex Cell MR Atto647N #NC.MR.100.AT). Briefly, blocking was carried out in the blocking buffer 60 min in a 37°C in a preheated humidity chamber. Primary antibodies were diluted in 80 µL of the provided diluent at 1/200 for both anti-p24 (mouse, from the NIH AIDS Reagents repository cat. ARP-4121 and ARP-3537) and anti-alpha tubulin (rabbit, Sigma cat. SAB4500087) and incubated overnight with gentle shaking at +4°C (in a home-made humid chamber), together with the anti-FLAG antibody (Goat, Novus Biologicals cat. NB600-344). Following this incubation, the primary antibody solution was decanted, and the wells were washed 2x 10 sec and 1x 15 min with 1X TBS-T under gentle shaking. The secondary antibodies from the kit and enzymes were diluted 1/80th instead of 1/40th to decrease background noise. Wells were then incubated for 1 hour at 37°C with secondary antibodies specific for the FLAG. Cells were finally stained with DAPI and imaging was done in PBS.

For live imaging of virion cores, 100000 TZM-BL cells expressing or not mscarlet-Trim69 were seeded in glass-bottom 35mm µ-Dish (IBIDI cat. 81156) in DMEM without phenol red (ThermoFisher cat. 31053028) supplemented with 7,5% serum. After 1 hour of incubation with VSVg-Env pseudotyped, Vpr-nG-IN virus, cells were stained with SiR-tubulin at 1µM supplemented with 10µM Verapamil, following the manufacturer’s instructions (Tebubio cat. SC002) for one additional hour prior to imaging. Images were acquired with the Confocal Zeiss LSM980-AiryScan 2 using a 63x/1.4 oil DIC Plan-Apochromat objective (Zeiss), 3x internal magnification at a frame size of 800 x 800 pixels and controlled by ZEN blue software. Live imaging was performed by using three laser sources: a 488 nm laser for exciting viruses, a 561 nm laser for exciting mscarlet-TRIM69 and a 638 nm laser for exciting SiR-tubulin. 2D acquisitions were carried out at a rate of one frame per 7,52 sec for 3-5min (total of 30 images). All acquired images were processed and visualized using Fiji software and virion cores tracking was analyzed with TrackMate plugin ^45^. Live imaging of mitochondria was performed using the PKmito RED (added to cell medium for 30min prior to acquisition at 37°C, 1/1000 from reconstitute stock as recommended by manufacturer, Tebubio cat. SC052). 2D images were acquired at a rate of one frame per 11,8 sec for 5min (total of 25 images). Two and three independent experiments were respectively analyzed for virions cores (8-9 cells per condition, between 283 and 329 particles analyzed) and mitochondria (18 cells per condition, between 4700 and 7500 particles analyzed) to determine the particle displacement, mean speed and the total distance traveled.

### Electron microscopy and immuno-gold EM

Ectopically transfected HEK293T cells were fixed for one hour with 4% paraformaldehyde in phosphate buffer (pH 7.6), washed and then infused with 2.3M sucrose for 2 hours (4°C). 90nm ultra-thin cryosections were generated at-110°C on a LEICA UCT cryoultramicrotome. Sections were retrieved with a 2%methylcellulose/2.3M sucrose mixture (1:1) and collected on formvar/carbon coated nickel grids. Sections were incubated with anti-Flag and anti-tubulin antibodies (Sigma cat. F7425 and T5168,). After extensive washing, grids were incubated with gold-conjugated goat-anti-rabbit IgG (Aurion, 10nm) and goat-anti-mouse IgG (Sigma, 6 nm). Grids were washed, post-fixed in 1% glutaraldehyde and rinsed. A contrasting step was performed by incubating grids on drops of uranyl 4%acetate/2%methycellulose mixture (1:10). The sections were imaged on a transmission electron microscope at 100 kV (JEOL 1011, Tokyo, Japan).

### siRNA-mediated silencing and RT-qPCR

Doxycycline-inducible TZM-BL cells were transfected with 40 nM of siRNA duplexes with Jetprime. Fresh medium with doxycycline was changed 2 hours after transfection. Cells were cultured for 24-48 hours before challenging or harvesting for the knock-down efficiency analysis. DYNC1H1 and BICD2 knockdowns were performed using the following siRNA duplexes: GAUCAAACAUGACGGAAUU and GCAACUGUCUCGUCAAAGA (J Virol. 2018 Oct 15; 92(20): e00358-18). siRNA duplex negative control was used as control (Eurogentec, cat. SR-CL000-005). To control the efficiency of silencing we used either WB (for BICD2) or RT-qPCR (DYNC1H1, as we could not identify a reliable antibody for WB analysis). In this case, total cellular RNA was extracted according to the manufacturer’s instructions (Macherey-Nagel NucleoSpin RNA cat. 740956) and reverse transcription was performed using random hexamers and oligo(dT) with the SuperScript III reverse transcriptase (Thermo Fisher cat. 18080). qPCRs were performed on a Biorad real-time PCR system (CFX Opus 96) using the FastStart universal SYBR green master mix (Roche Diagnostics cat. 4913914001). The primers used for mRNA quantification of DYNC1H1 (forward: 5’ GGTTTGGCTTCAGTATCAGTG 3’ and reverse: 5’ CTGGTCCAAATAAGGAAGGCC 3’) were standardized against non-targeting siRNA control, with actin (forward 5’ TTTTCACGGTTGGCCTTAGG 3’ and reverse: 5’ AAGATCTGGCACCACACCTTCT 3’) levels as an internal control.

### Cell mortality and cell cycle analyses

Cell mortality was assessed in Trim69 stable cells either in a cycling (TZM-BL), or in a growth-arrested status (THP-1-PMA, mirroring macrophage-like conditions). Cells were harvested at different times post Trim69 induction, they were stained either with PE Annexin V Apoptosis Detection Kit I (BD Biosciences cat. 559763) for apoptosis analysis or with propidium iodide (PI, SIGMA cat. P4864-10ML) for cell cycle analysis using flow cytometry. To determine the distribution of cell cycle phases, cells were fixed in 70% ethanol, then resuspended in 500ml of PBS 1X containing PI and RNaseA at final concentrations of 0.5mg/ml and 100mg/ml, respectively, for 30min prior to analysis by flow cytometry. Cells in G1, S and G2/M phases can be distinguished thanks to the ability of PI to intercalate proportionally to the amount of DNA present in the cell. The mitotic index of cell cultures was determined by confocal microscopy imaging of cells stained with an anti phospho-H3 antibody (Cell Signaling cat. 53348T) that labels the nuclei of all cells undergoing mitosis. The proportion of phospho-H3-positive cells among DAPI-positive nuclei was counted visually in pictures taken at 20x magnification.

### Tubulin preparation

Tubulin was purified from mouse brains as described before ^46^ using the polymerization/depolymerization method. Animal care and use for this study were performed in accordance with the recommendations of the European Community (2010/63/UE) for the care and use of laboratory animals. Wild type mice were sacrificed by cervical elongation and their brains were removed rapidly and homogenized (2 ml of buffer/1 g of brain tissue) in lysis buffer: BRB80 (BRB80: 80mM PIPES pH 6.9, 1mM EGTA, 1mM MgCl2) supplemented with 1 mM 2-mercaptoethanol, 1 mM PMSF, and protease inhibitors. The lysate was clarified by ultracentrifugation at 180,000 x g at 4°C for 30 min (Beckman Optima, TLA55 rotor). The supernatant was supplemented with 1mM GTP (Sigma) and 30% glycerol and incubated at 30°C for 30 min (1st microtubule polymerization). The sample was then spun down at 180,000 x g at 30°C for 30 min, supernatant removed, and the pelleted microtubules were resuspended in BRB80 (1/20th of the volume of the clarified lysate). After pipetting up and down for 20 min on ice to depolymerize microtubules, the sample was spun down at 180,000 x g at 4°C for 30 min. The supernatant (containing soluble tubulin) was supplemented with 1 mM GTP, 0.5 M PIPES-KOH pH 6.8 and 30% glycerol, and incubated at 30°C for 30 minutes (second polymerization, in the presence of high PIPES leading to the exclusion of microtubule-associated proteins from the microtubules). The sample was spun down at 180,000 x g at 30°C for 30 minutes, and the microtubule pellet was depolymerized in BRB80. One more polymerization/depolymerization (in low PIPES conditions) cycle was performed to remove traces of 0.5 M PIPES buffer. Pure tubulin was then used to prepare Taxol-stabilized microtubules used in the TIRF assay by incubating 4 mg/ml tubulin with 4 mM MgCl2, 5% DMSO, and 1 mM GTP for 30 min at 37°C. The polymerized microtubules were then diluted in BRB80T (BRB80 supplemented with 10 µM Taxol) and centrifuged for 30 min at 18,000 x g. After centrifugation, the microtubules were resuspended and kept in BRB80T at room temperature and used within 2 weeks.

### Subtilisin-treated microtubules

Taxol-stabilized microtubules were prepared as described above. Subtilisin (Sigma, cat. P8038) was added to polymerized microtubules at an enzyme:tubulin ratio of 1:10 and incubated for 30 min at 30°C. The reaction was stopped by the addition of 1 mM PMSF (Sigma, cat. 78830) and incubated for 10 min at room temperature. The modified microtubules were then centrifuged at 20,000 x g for 20 min at room temperature after which they were resuspended in BRB80T and used within 2 weeks. Trim69-wt was diluted in AB buffer to the desired concentration and incubated on the modified microtubules for 3 min.

### Cell lysate preparation

Cell lysates overexpressing YFP-tagged proteins were obtained as described before ^27^ ^28^. Briefly, HEK293 cells were transfected using the jetPEI (Polyplus Transfection), according to manufacturer’s instructions. After 24 to 48 hours, cells were collected and resuspended in lysis buffer (BRB80 supplemented with 0.05% Triton X-100 and protease inhibitors) and lysed on ice for 10 minutes by pipetting up and down. Lysates were then sonicated for 3 sec and spun down at 95,000 x g for 25 min at 4°C (Beckman Optima, TLA55 rotor). The supernatant was removed, snap frozen, and kept at-80°C for up to 2 months.

### Experimental chamber preparation

For TIRF microscopy assays, experimental chambers were assembled by melting thin strips of parafilm in between two glass coverslips (22x22 mm, VWR #630-2186 and 18x18 mm, VWR #630-2185) cleaned by subsequent 15 min sonication steps in 3M KOH, osmosis water, and absolute ethanol (VWR #20821.310). The glass coverslips were kept in replenished absolute ethanol at room temperature for up to 2 months. The experimental chambers were incubated with 8 µM rigor kinesin for 5 min, followed by an incubation with 1% Pluronic (F127 in PBS, #P2443, Sigma) for at least 30 min. Microtubules, visualized by total interference reflection microscopy (IRM), were flushed into the chamber and allowed to adhere to the rigor kinesin. Unbound microtubules were washed away with BRB80T and chambers were pre-incubated with TIRF assay buffer AB (BRB80, 10 mM dithiothreitol, 5,6 µg/ml casein, 10 µM Taxol, 1 mM Mg-ATP, 20 mM D-glucose, 0.22 mg/ml glucose oxidase and 20 µg/ml catalase) prior to experiments. Finally, cell lysates were diluted in buffer AB to the desired concentration and added to the measurement chamber.

### TIRF microscopy

Total internal reflection fluorescent (TIRF) microscopy assays were performed on an inverted microscope (Nikon Eclipse Ti) with FRAP-TIRF module (Ilas2), equipped with motorized XY-stage, perfect focus system, Nikon Apo TIRF 100x oil immersion objective (NA 1.49), and EMCCD Evolve 512 camera. Microtubules were visualized using interference reflection microscopy (IRM) and fluorescent YFP-tagged proteins by using a FITC filter cube.

### Analysis of cell-lysate-based IRM-TIRF assays

Trim69 (WT or SPRY-7A) density on microtubules was measured by drawing a line along the microtubules on the IRM image using ImageJ (version 2.14.0/1.54f) and transferring this line to the TIRF image to measure the mean intensity of Trim69 along the MT. Background subtraction was done for each microtubule by subtracting the mean intensity of an identical area adjacent to the microtubule where no microtubule was present.

### Fluorescence recovery after photobleaching (FRAP) experiments

In vitro FRAP experiments were carried out using the FRAP module (Ilas2) on the TIRF microscope. YFP-Trim69-wt lysate (1000x diluted in AB) was incubated on Taxol-stabilized microtubules for at least 3 min to establish stable binding to the microtubules. Microtubules were imaged for 1 min pre-FRAP to establish the baseline density of the Trim69 signal. FRAP was then performed on a rectangular region along the MT after which the MT was imaged for 6.5 min (5 sec framerate). The post-FRAP density was normalized to the pre-FRAP density at frame 1. For the non-FRAP curve, Trim69 signal on MTs that were not subjected to photobleaching were analyzed and normalized by the same method.

For FRAP experiments in cells, TZM-BL cells were transfected with mscarlet-TRIM69 24 hours before imaging. Photoactivation were performed using Confocal Zeiss LSM980-AiryScan 2 using a 63x/1.4 oil DIC Plan-Apochromat objective (Zeiss), 3x internal magnification at a frame size of 256 x 256 pixels and controlled by ZEN blue software. Two different regions of filamentous TRIM69 were targeted for either photobleaching or control (without photobleaching). Pre- and post-photobleaching time frames were acquired with 561 nm laser with 9 sec intervals for around 4 minutes. For acquisition, the laser power used was 50% and the gain was adjusted to 800. Cells were imaged for 45sec before photobleaching (prebleaching intensity). Fluorescence recovery of the two regions of interests (ROI) was analyzed with the Zen blue 2.3 software which allows to extract fluorescence intensity values over time.

### Structure modeling

The model was generated using AlphaFold 2.0, following by an ESPript analysis of AlphaFold models ^23,24^, PRABI Lyon-Gerland Facility Service). Only the first four models with an AlphaFold score exceeding 0.45 were considered. Critical hydrogen bonds (H-bonds) between the SPRY domain of Trim69 and the C-terminal tail of α- or β-Tub (Tubb2b) were determined using ChimeraX ^47^, with a distance tolerance of 0.4 Å and an angle tolerance of 20°. Alpha Fold 3.0 webservice was used to analyze isolated SPRY domain of Trim69 and the C-terminal regions of α- and β-tubulin (AlphaFold webserver, ^26^).

### Trim69 sequence analyses

The mammalian TRIM69 amino acid sequences were retrieved using OrthoMaM (Orthologous Mammalian Markers, ^48^) v12a, using human TRIM69 (ENSG00000185880) as query. A final number of 158 high-quality sequences were retained and aligned using PRANK v.170427 ^49^. The basic residues important for human TRIM69 antiviral functions were located on the alignment and the sequence logo of the surrounding region was generated using WebLogo3 (https://weblogo.threeplusone.com/).

### Softwares

Confocal microscopy software: Fiji (version 2.0.0), Zen (version 2.3, Zeiss) and Imaris 9.2.0 (Oxford Instruments Group). TIRF microscopy software: Metamorph (version 7.10.5.476). WB: Image Lab Touch (version 2.0.0.27, Chemidoc Imaging System from Bio-Rad). Flow cytometry: FlowJo (version X, BD). Protein structure modeling/analysis: ChimeraX (version 1.6.1) and AlphaFold 2.0 and 3.0. Sequence analysis: Geneious Prime 2024.0.3. Statistics and graphs: Graphpad Prism8 (8.4.3, Graphpad software, LLC) and Excel (16.16.3, Microsoft).

### Statistics

Statistical analyses were performed using GraphPad Prism8, as described in each legend.

## Data availability

Data generated in this study are included in the Paper. Source data are provided.

The mammalian TRIM69 amino acid alignment is openly available on Figshare in FASTA format, along with a PDF visualization of the alignment obtained with ESPript v3 (https://espript.ibcp.fr) ^24^. These files can be accessed at https://doi.org/10.6084/m9.figshare.28330211.v1 and https://doi.org/10.6084/m9.figshare.28330208.v1.

## Acknowledgments

C.V. is the recipient of a PhD fellowship from the ANRS|MIE (AO-2021-1, ECTZ162940) and the laboratory is supported by grants from the ANRS|MIE (AO-2021-1, ECTZ160109 and ANRS0644), Sidaction (AAP34-1 – 13904) and ANR (ANR-24-CE15-2986, the latter to A.C, F.F. and M.M.M.). The Janke lab is financed by the ERC Synergy TubulinCode to C.J. A.Ch. is the recipient of a fellowship from the ENS-Lyon.

We thank Olivier Reynard (CIRI, Lyon) for sharing material and the authors that provided plasmids through Addgene. We are indebted to Francis Ashwanth (Florida State University, USA) for sharing material and technical insights on viral cores imaging. We acknowledge the contributions of the SFR Biosciences (Universite Claude Bernard Lyon 1, CNRS UAR3444, Inserm US8, ENS de Lyon) (LYMIC-PLATIM) and of the center AniRA in Lyon of the CELPHEDIA Infrastructure (http://www.celphedia.eu/). We also thank: Z. Lansky/M. Braun lab members for helpful discussions (BIOCEV Institute of Biotechnology, Czech Republic), S. Ramakrishnan, R. Onclercq-Delic, M. Högner from the Janke lab for technical help. We also acknowledge K. Bellul, C. Jouhanneau, V. Dangles-Marie and H. Gautier from the Institut Curie animal facility, and M.-N. Soler and L. Besse from the Multimodal Imaging Center of the Institut Curie (CNRS UAR2016/ Inserm US43/ Institut Curie / Université Paris-Saclay).

A.C., L.E., C.J., F.F. and M.M.M. are researchers of the Centre National de la Recherche Scientifique (CNRS). The funders had no role in study design, data collection and analysis, decision to publish, or preparation of the manuscript.

## Author contributions

C.V. and X.N.N. designed, performed and analyzed *in cellulo* experiments; V.S. and A.K. performed and analyzed TIRF experiments; V.H. contributed new reagents; A.Ch. and L.E. performed phylogenetic analyses; J.B.G. and P.R. performed electron microscopy analyses, F.F. performed structural analyses; C.J. and M.M.M. analyzed MT experiments; F.F., M.M.M. and A.C. designed the study, analyzed data, acquired funding and wrote the paper.

## Ethics declarations

Competing interests. The authors declare no competing interests.

## Extended Data

**Extended Data 1. Complete channel presentation of the confocal microscopy analysis of Fig 1C and cell toxicity, cell cycle and mitotic index analyses of cells expressing Trim69. A)** The figure presents the individual and merged IF channels for the IF presented in Fig 1C. **B)** Immuno-gold electron microscopy of THP-PMA cells with anti-tubulin or anti-Trim69 specific antibodies. **C)** Cycling TZM-BL and growth-arrested macrophage-like THP-PMA cells stably expressing doxycycline-inducible Trim69, or an identical vector devoid of Trim69 were treated with dox dox (1.5 μg/mL for 24-48 hours). Cells were analyzed by flow cytometry at the indicated time following AnnexinV staining to determine the percentage of apoptotic cells. **D)** As in C, but cells were first fixed and then stained with propidium iodide prior to flow cytometry analysis to determine whether Trim69 altered the cell cycle (a representative result of cells at day 4 is provided). **E)** As above, but cells were analyzed by confocal microscopy upon staining with the phosphorylated form of histone 3 (H3-P), present in cells undergoing mitosis. Cells were visually scored for positivity on large fields. The number of H3-P positive cells within a culture yields a measure of the mitotic index of a cell population, i.e. the number of cells undergoing mitosis in an asynchronous cell population. Given that they do not cycle, this analysis was not carried out in THP-PMA cells. No statistically significant differences were observed between control and Trim69 cells following individual Student t tests in the different experiments.

**Extended Data 2. Structural modeling. A)** Alignment and analyses of secondary structures extracted from Alphafold 2.0 best-ranked models in two conditions, as indicated. The α and ɳ helices are displayed as squiggles; β-strands as arrows; turns as TT letters. The secondary structures are colored according to the AlphaFold confidence score in the accuracy of the prediction (pLDDT - Predicted Local Distance Difference Test): Blue (90-100 pLDDT) represents very high confidence; Cyan (70-90 pLDDT) high confidence; Yellow (50-70 pLDDT) moderate confidence; and Orange/Red (<50 pLDDT) low confidence. In the middle: Sequence alignment of Tubb2b proteins from different species: red shading=strictly identical residues; yellow shading= similar residues. The relative accessibility (acc) of each residue is represented by colored boxes: blue for accessible, cyan for intermediate accessibility, and white for buried. The hydropathic (hyb) character is also represented by colored squares: pink for hydrophobic, grey for intermediate and cyan for hydrophilic residues. At the bottom: intramolecular contacts present in each model are indicated by the letter of the chain. In this case, the analysis was performed using β-tubulin (chain C) as the query and Trim 69 (chain A) and α-tubulin (chain B) as interactors. Red lettering indicates a putative binding distance of < 3.2 Å, while black lettering signifies distances between 3.2 and 4 Å. **B)** Resides involved in the interaction. Underlined indicates residues present in a flexible loop of the SPRY domain of Trim69 and its putative interactor amino acid on Tubb2b. **C)** as in B on the 3D structures of Trim69 (SPRY domain, grey) and Tubb2b (C-terminal tail, cyan). **D)** Comparison and overlay between the SPRY domains of Trim69 and of Trim5α (violet) with respect to the fitting of the C-terminal tail of β-Tub. V1 to V3 indicate the variable loops of Trim5α important for its binding to retroviral capsids.

**Extended Data 3. Sequence variations in the C-terminal tails of human tubulin isotypes.** Sequence alignment of the α- and β-tubulin isotypes coded in the human genome. Residues potentially involved in the interaction with the SPRY domain are highlighted in color.

**Extended Data 4. Binding of Trim69 to microtubules in vitro and conserved ability of the SPRY-7A mutant to undergo Trim69:Trim69 interactions. A and B)** Representative TIRF image of wild type and SPRY-7A YFP-Trim69-expressing lysates (bottom panels) diluted 100x (left) or 10x (right) in assay buffer and incubated for 3 min on label-free Taxol-stabilized microtubules (top panels, visualized by IRM). Scalebars: 2 µm. **C)** Density of wild type YFP-Trim69 (red) and mutant YFP-Trim69-SPRY-7A (grey) on Taxol-stabilized microtubules at different lysate dilutions. Wild type Trim69 density at 100x lysate dilution was 731 ± 265, and at 10x: 2980 ± 1161 (n=57, 58 microtubules in 5 independent experiments, respectively). Mutant Trim69-SPRY-7A density at 100x lysate dilution was 2 ± 15, and at 10x: 4 ± 49 (n=43, 34 microtubules in 10, 8 independent experiments). *, p< 0.05 following a two-sided Student t-test. **D and E**) HEK293T cells expressing two versions of Trim69 (YFP-Trim69 and Flag-Trim69) distinguishable by size upon WB and also by confocal microscopy were used to assess the ability of the SPRY-7A mutant to undergo Trim69:Trim69 interactions by co-immunoprecipitation (D) and confocal microscopy (E). Representative results are shown in the different panels.

**Extended Data 5. The basic patch of the SPRY domain is conserved across mammalian Trim69.** Logo plots illustrating the conservation of residues in the regions surrounding the seven sites predicted to interact with β-tubulin. For each site, the human residue is highlighted in red.

**Extended Data 6. Qualitative analysis of the distribution of endogenous EB1 in TZM-BL cells.** TZM-BL cells expressing or not Trim69 were analyzed by confocal microscopy using the indicated antibodies. Representative pictures are shown.

**Extended Data 7. Complete channel presentation for the confocal microscopy analysis of Fig 4A, and distribution of EB1 in cells treated with Taxol.** Individual and merged-panels for Fig 4A are presented here along with additional measurements of EB1 comets in control and Trim69-expressing cells shown in Fig. 4A. These analyses indicate no change in the average length of comets per cell, nor in their individual length. The bottom pictures present the analysis of the EB1 distribution in control THP-PMA cells treated with Taxol dox (20 μM for 24 hours). Given the extensive spread of EB1 in this condition, the quantification of EB1 comets was not possible. Ns, non-significant following an unpaired, two-tailed, Student t test.

**Extended Data 8. Complete channel presentation for the PLA experiment of** Fig. 4B **and additional analyses indicating that Trim69 decreases the number of viral cores that dock on MTs. A)** Cells were challenged with HIV-1 virus pseudotyped with JR-FL and incorporating a Vpr-INmNeonGreen for two hours prior to confocal analysis of the number of cells presenting HIV-1 viral cores (green dots, the Trim69 signal is saturated to clearly distinguish positive cells). The graph presents AVG and SEM obtained from the analysis of independent large field pictures (1 dot=1 picture); three independent experiments with an overall total number of cells analyzed comprised between 119 and 180. B) The figure presents the separated channels for the PLA analysis presented in Fig. 4B.

**Extended Data 9. Effect of Ciliobrevin D on the antiviral effects of Trim69 and efficiency of siRNA-mediated silencing of dynein motor subunits. A)** TZM-BL cells expressing or not Trim69 were challenged with GFP-coding and JR-FL Env-bearing single round of infection-competent HIV-1 virus in the presence or absence of the dynein motor complex inhibitor Ciliobrevin D (20 μM for 1 hour prior to infection). The extent of infection was measured by flow cytometry two to three days later. **B)** Silencing efficiency of the dynein components and of the experiment presented in Fig 5A was determined by either RT-qPCR (DYNC1H1), or WB (BICD2) in 3 independent experiments and representative panels, respectively. Ns and *: non-statistically significant and p<0.05 according to an ordinary one-way Anova, Tukey’s multiple comparison test (A), or following a Student t-test (unpaired, two-tailed) between the indicated control and specific siRNA-treated sample pairs (B).

**Extended Data 10. Redistribution of endogenous DYNLL1 and complete channel presentation for the confocal microscopy analysis of** Fig. 5C and 5D. TZM-BL cells expressing or not Trim69 were both stimulated with dox prior to staining with antibodies specific for endogenous DYNLL1. Representative results are shown here. The rest of the figure presents the separated channels for the confocal microscopy analysis presented in Fig. 5D.

**Extended Data 11. Lack of redistribution of the dynein and dynactin complexes in cells treated with Taxol.** Control cells expressing the indicated components of the dynein and dynactin complexes were treated with the MT-stabilizing compound Taxol (20 μM for 24 hours) prior to confocal microscopy. Representative pictures are shown here and can be compared to control, untreated cells in Fig 5D or in the Extended Data 10 (n=2).

**Extended Data 12. Trim69 inhibits the migration of dynein-dependent cargoes: HIV-1 virion cores on MTs, and red-mitotracker-labeled mitochondria. A and B)** Cells were stained for 30 min with PKmito RED to label mitochondria, prior to live imaging. Images were taken every 11.8 sec for 3-5 minutes. Individual mitochondrial tracks were then reconstructed with TrackMate plugin in FIJI and measured. The panels present typical images obtained, while the graphs present the total distance travelled and the mean speed of individual mitochondria (between 4700 and 7500 individual trajectories per condition, measured in 7 to 10 individual cells. *, p<0.05 following an unpaired, two-tailed Student t-test between the indicated conditions. **C)** Large view of cells expressing Trim69 or not (as indicated) and containing the virion cores displayed in Fig. 7A.

