## Supplemental Figures for "Microtubule remodeling by the innate immune factor Trim69 compromises dynein-dependent migration of HIV virion cores towards the nucleus"

*Extended data 1*

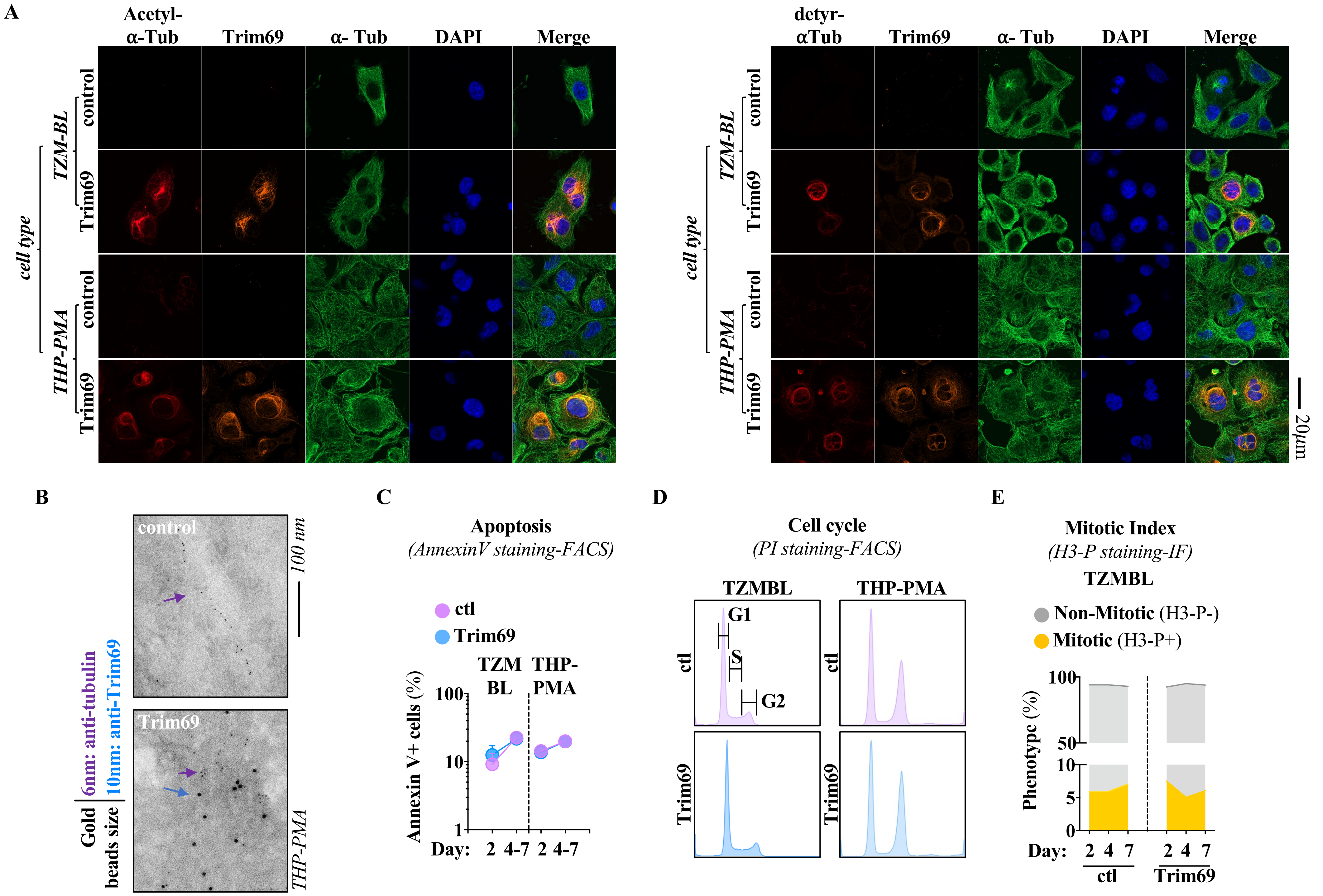

Extended data 2

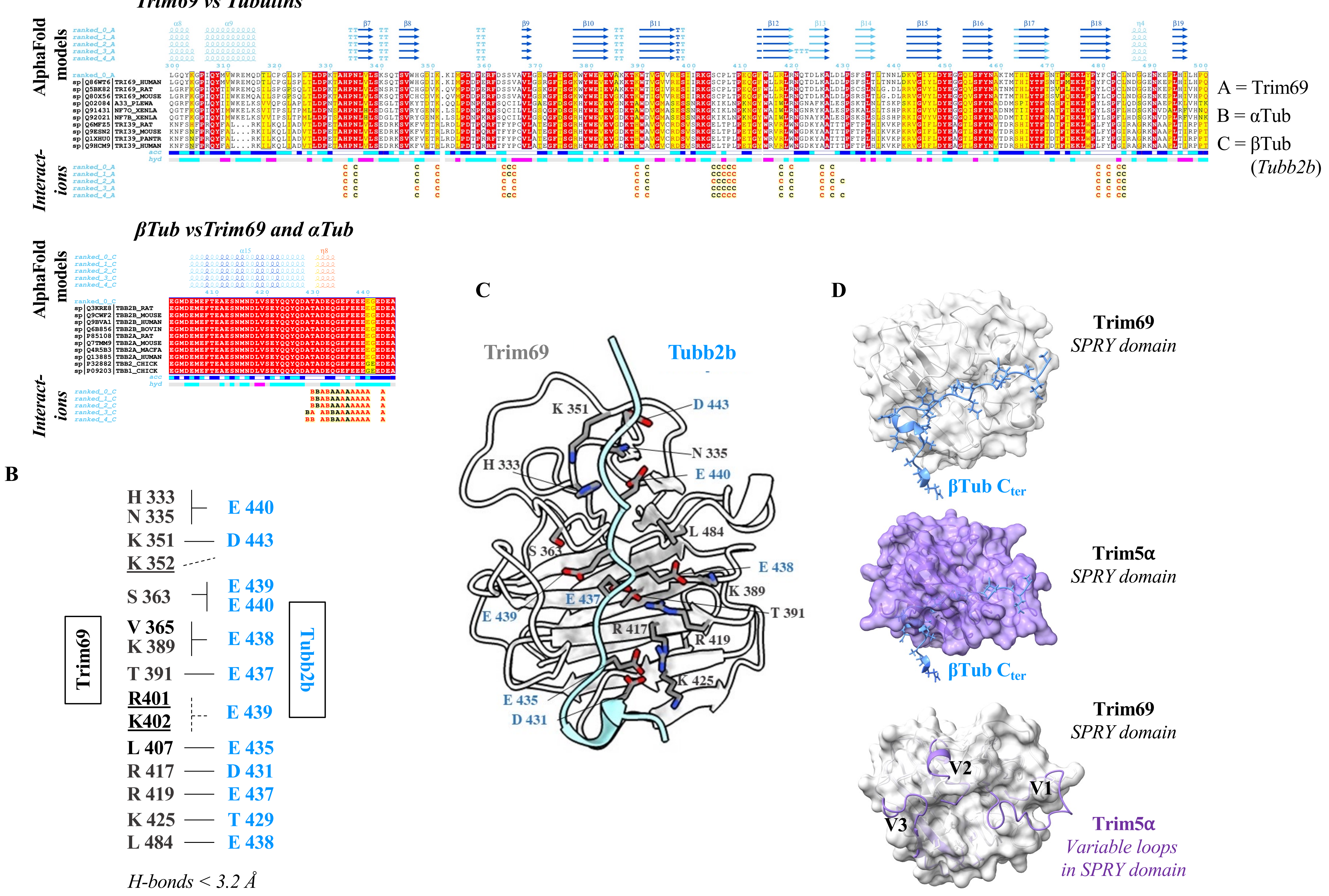

*Extended data 3*

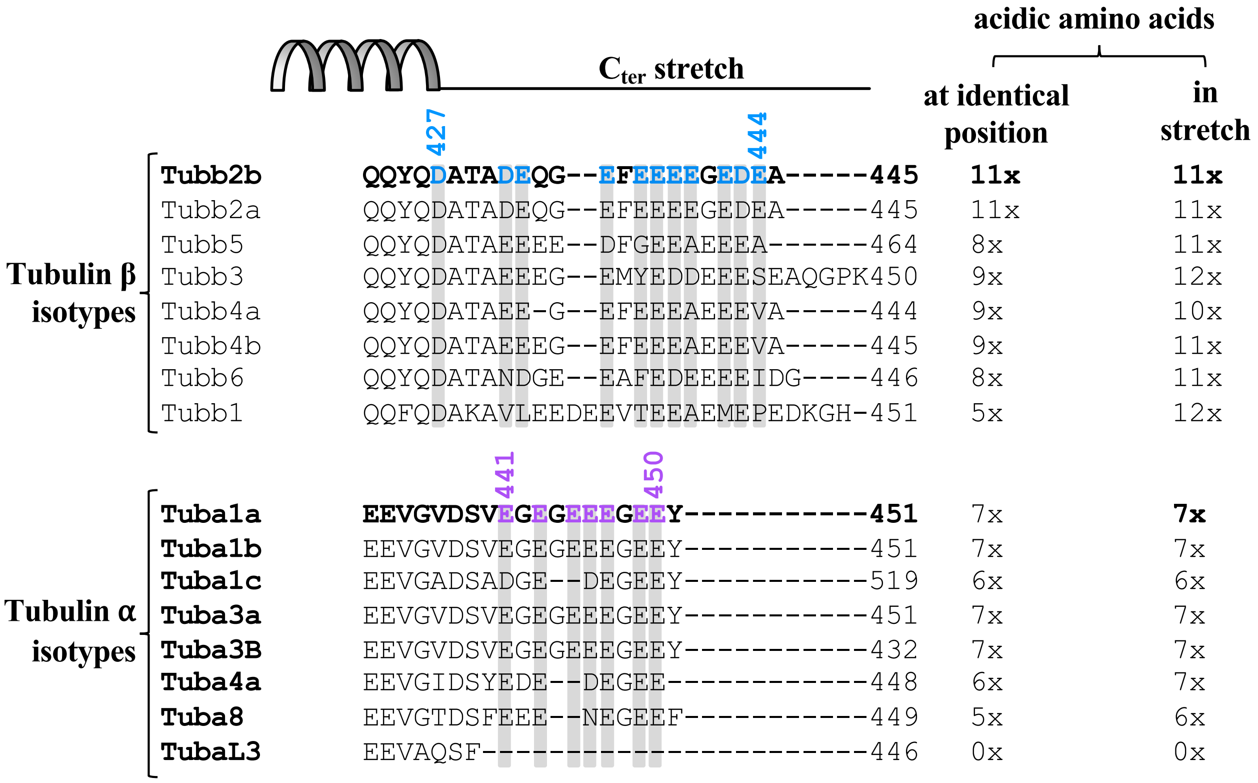

*Extended data 4*

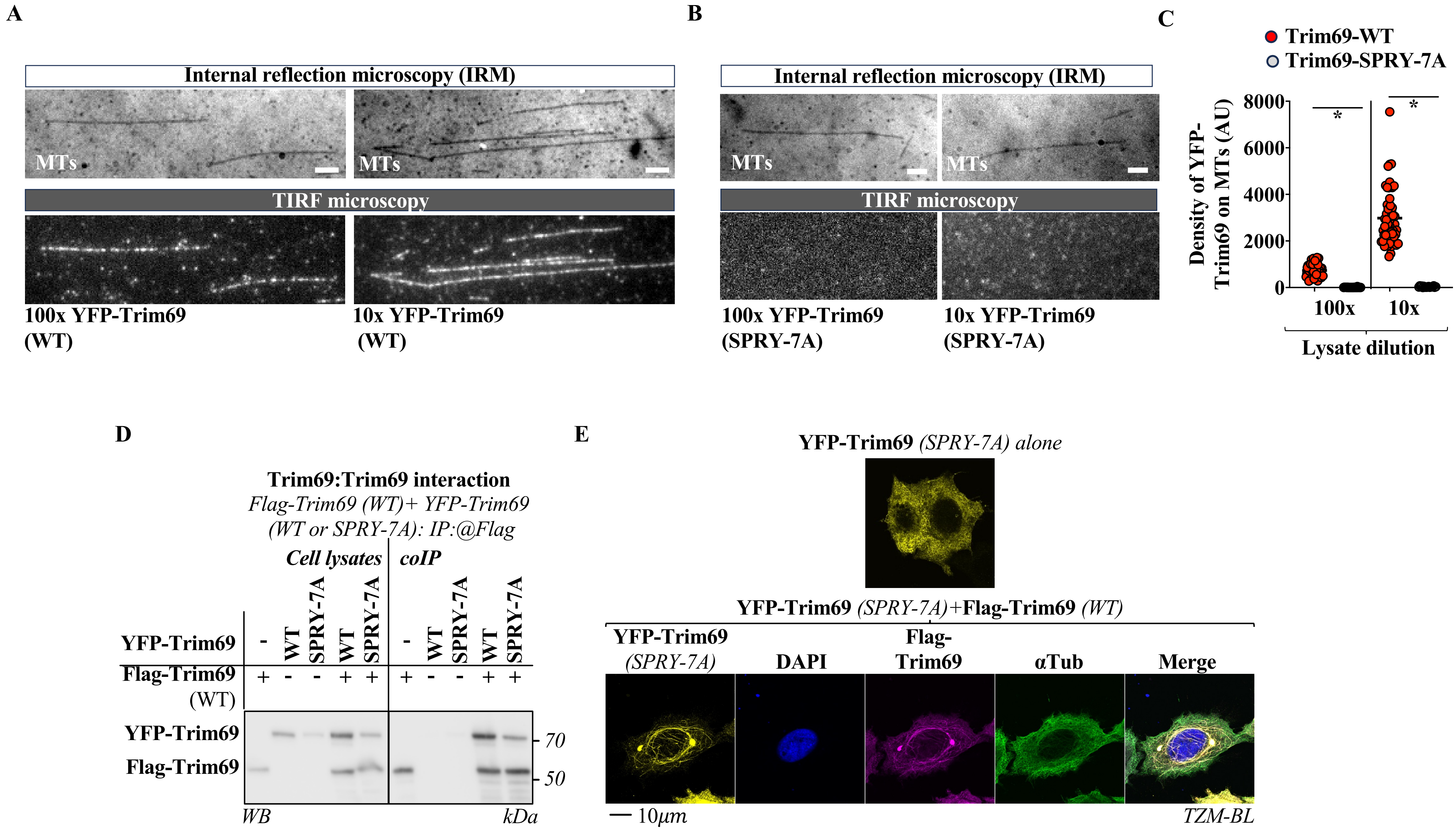

*Extended data 5*

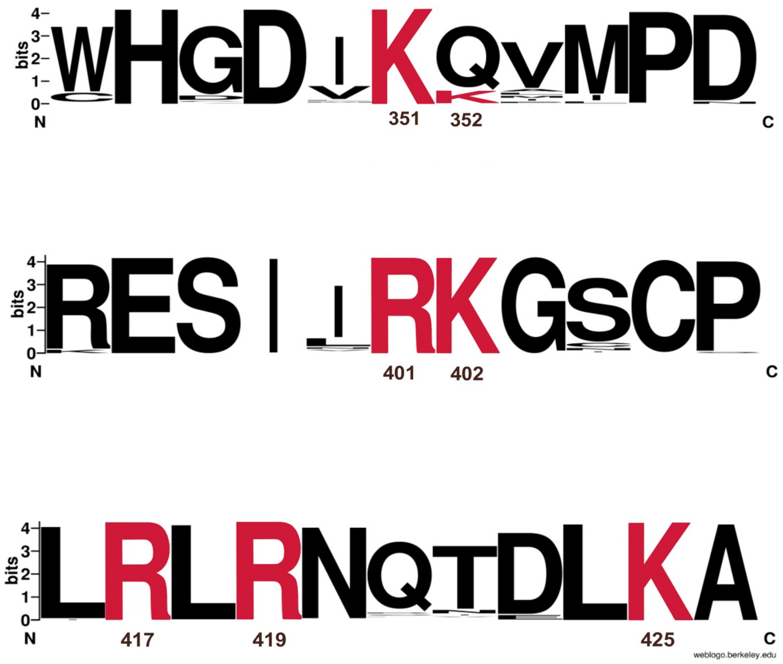

*Extended data 6*

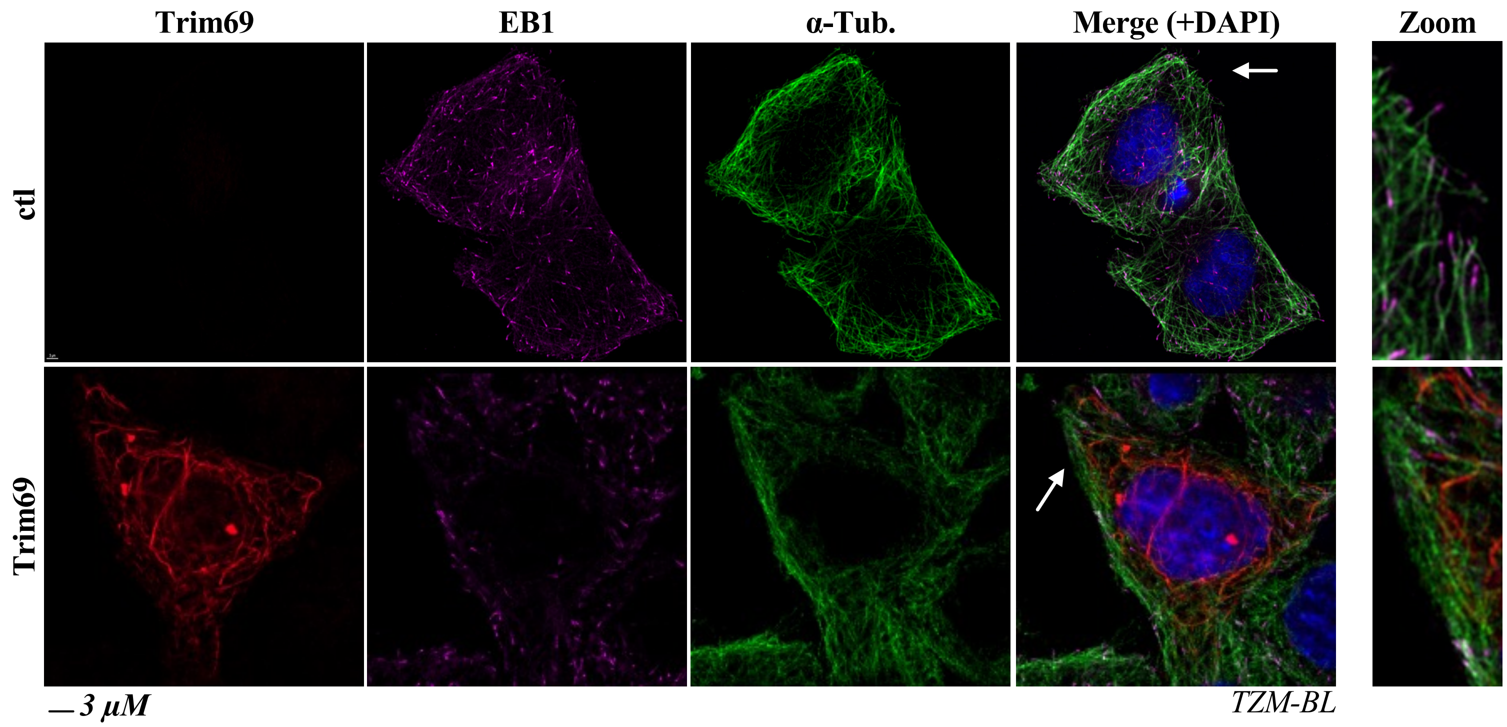

*Extended data 7*

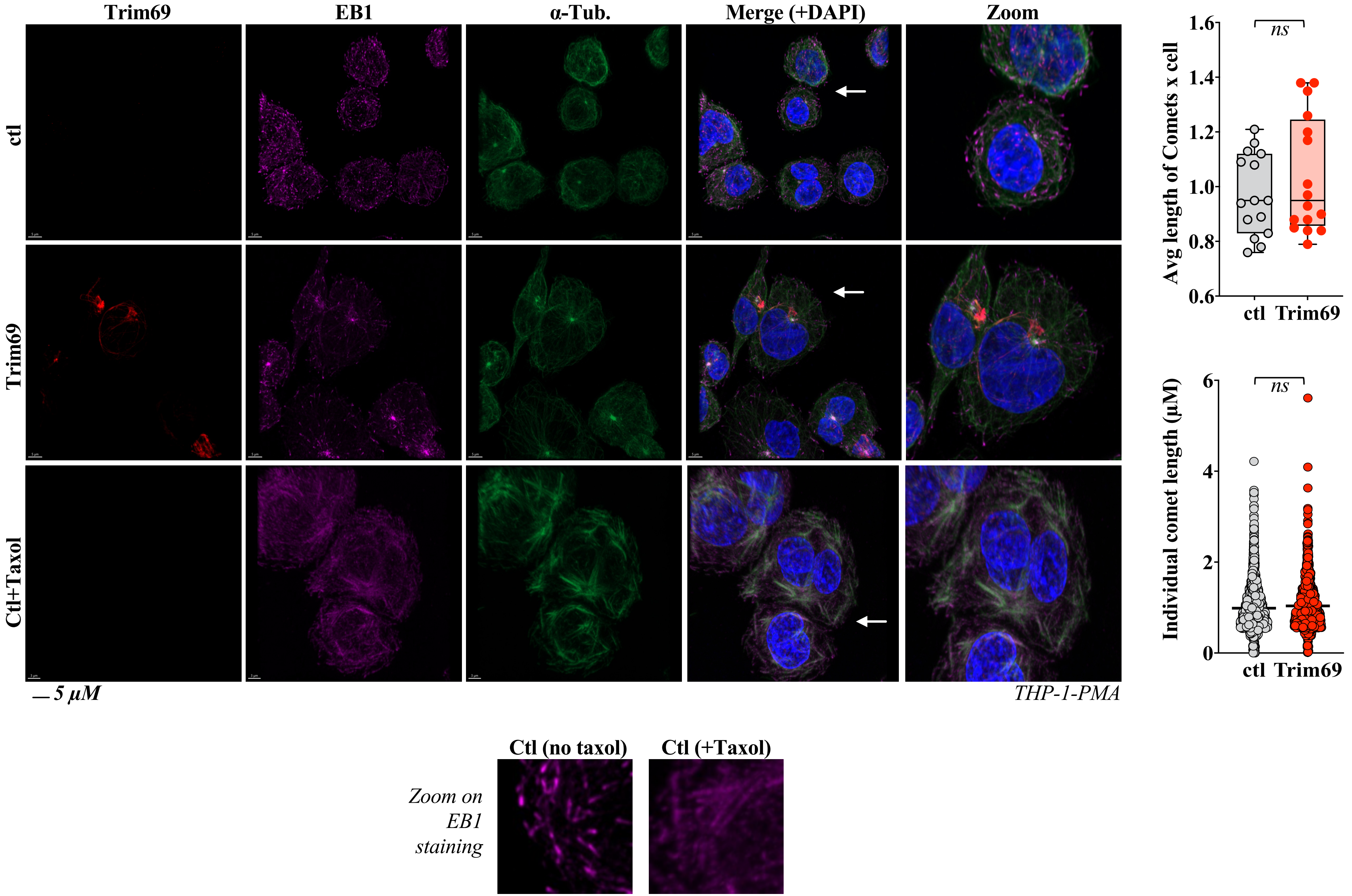

*Extended data 8*

**A**

**Proportion of cells with HIV-1 capsids** ***(Vpr-INmNG)***
*(1 dot=1 picture)*

**Example: HIV-1 capsids** ***(Vpr-INmNG)***
**+DAPI + Trim69 when indicated**
**(saturated)**

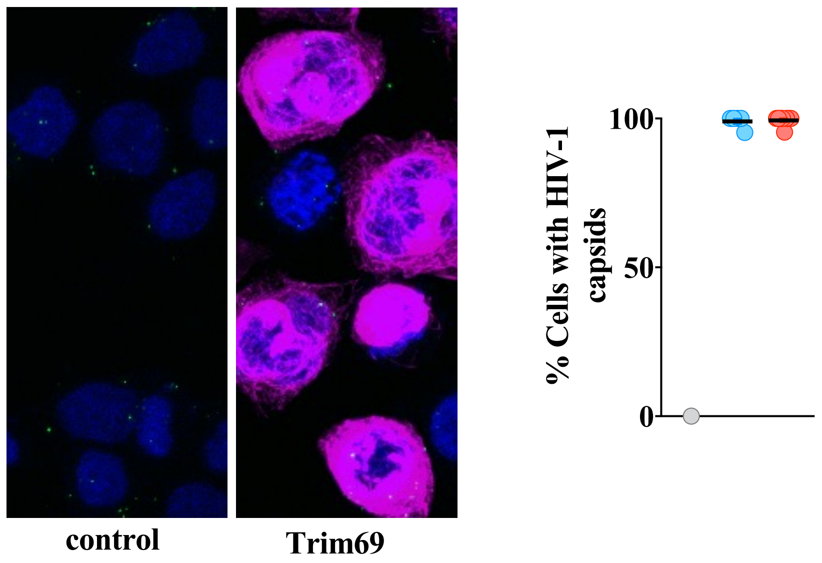


**B**

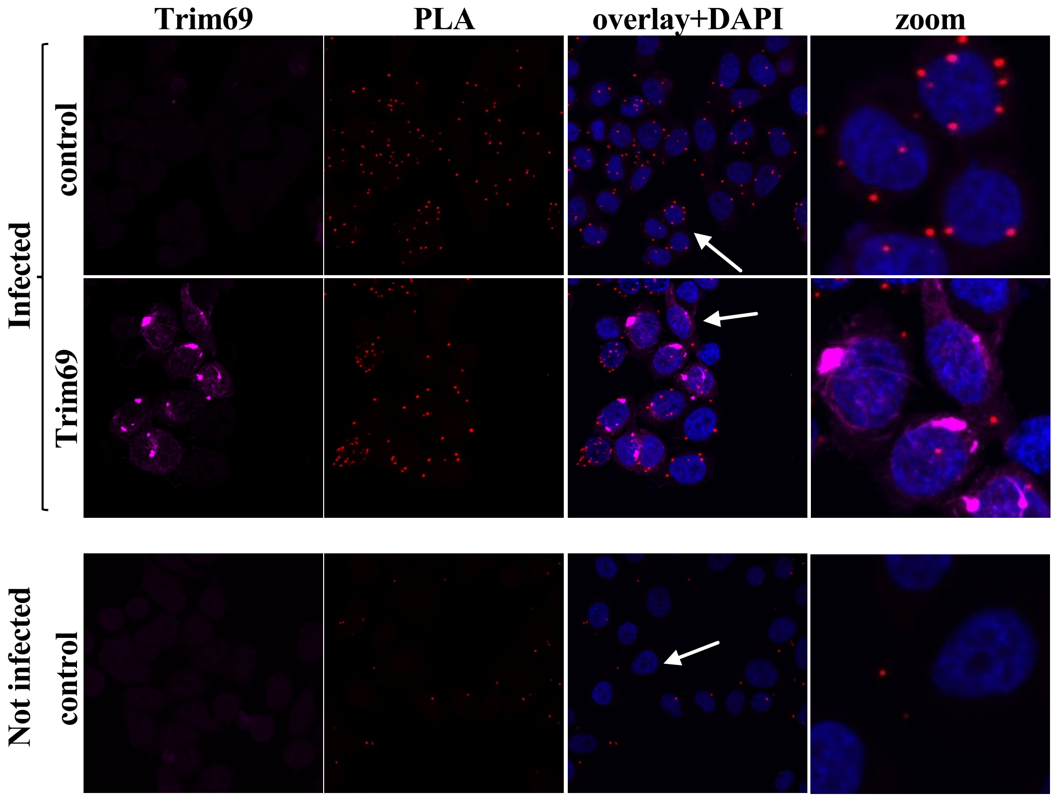

*Extended data 9*

**A**

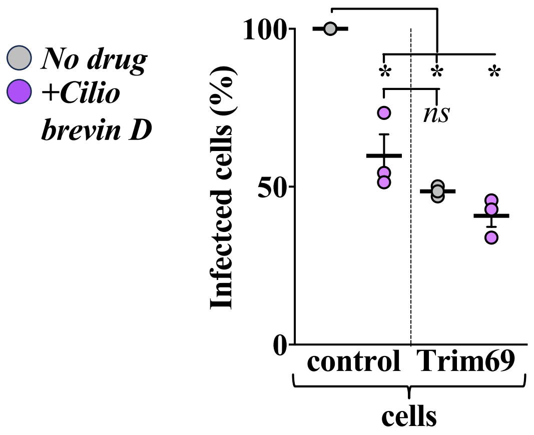


**B**

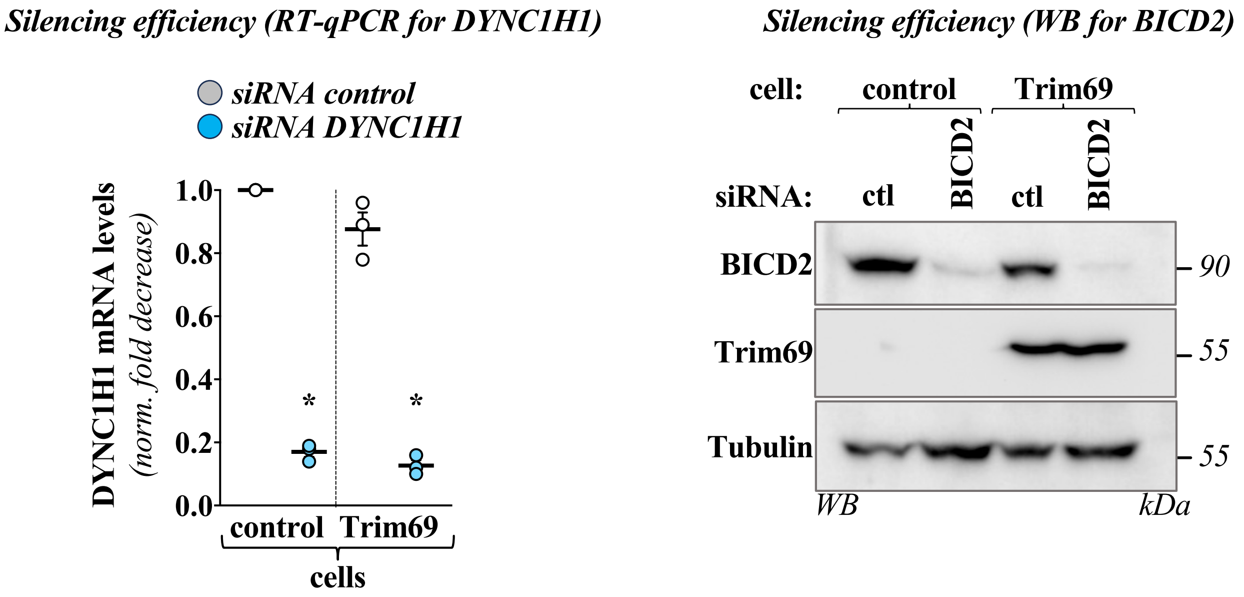

*Extended data 10*

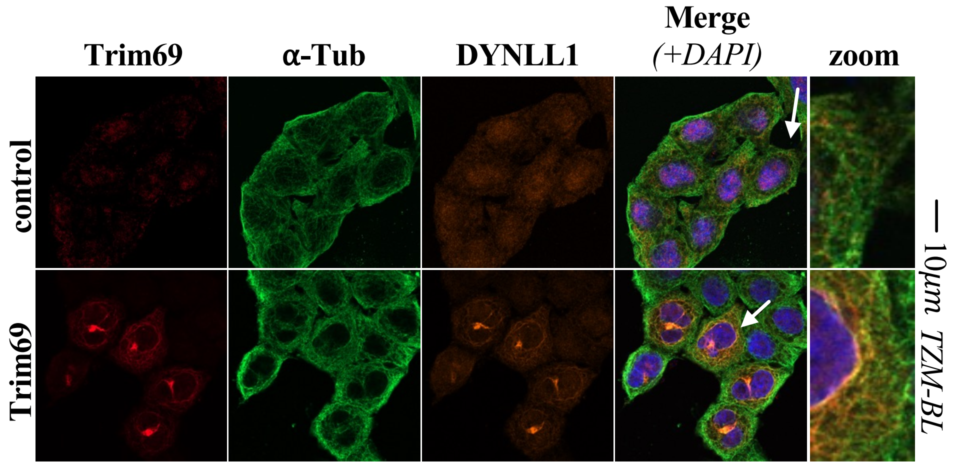

*Extended data 10, continued*

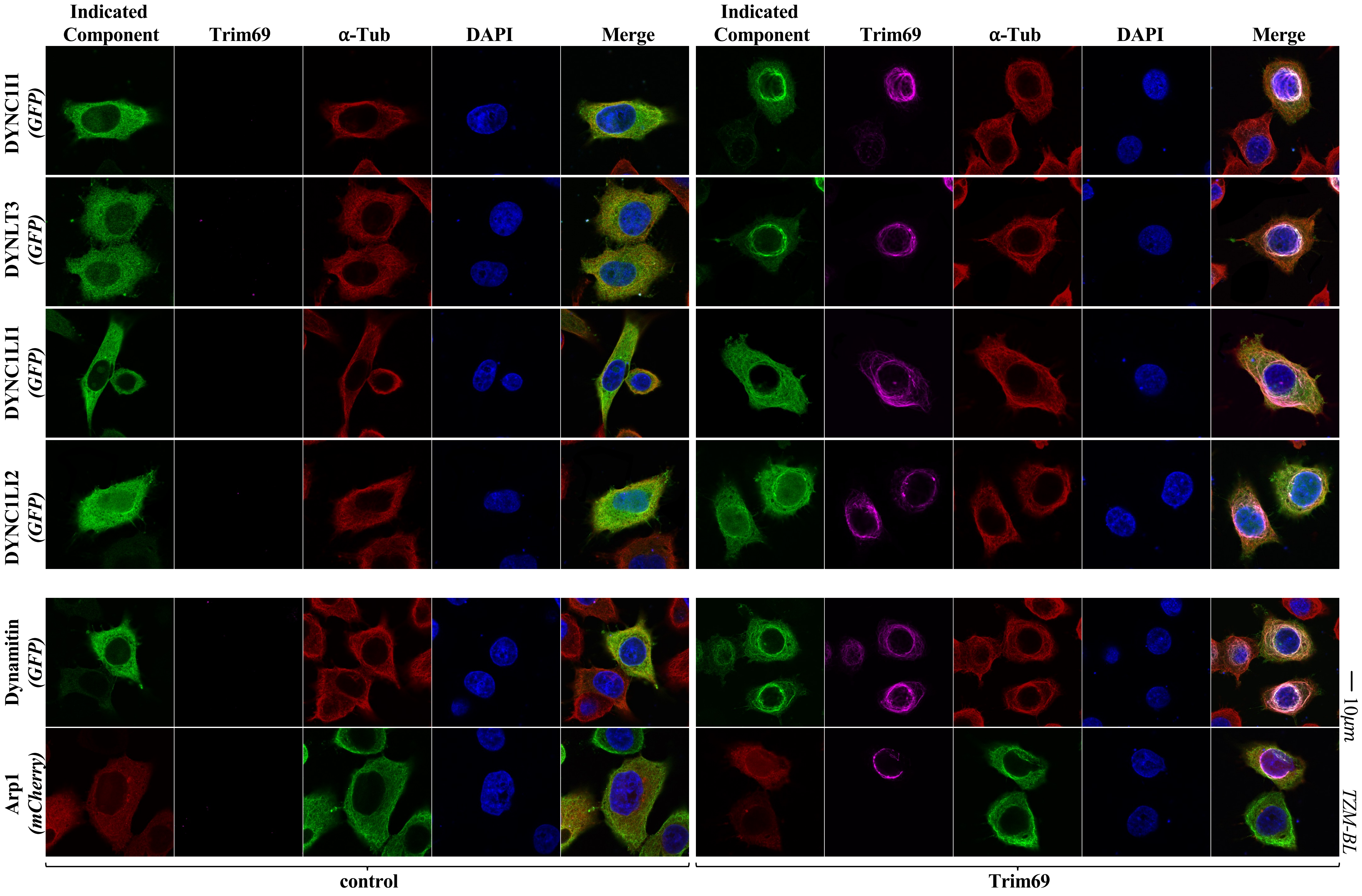

*Extended data 11*

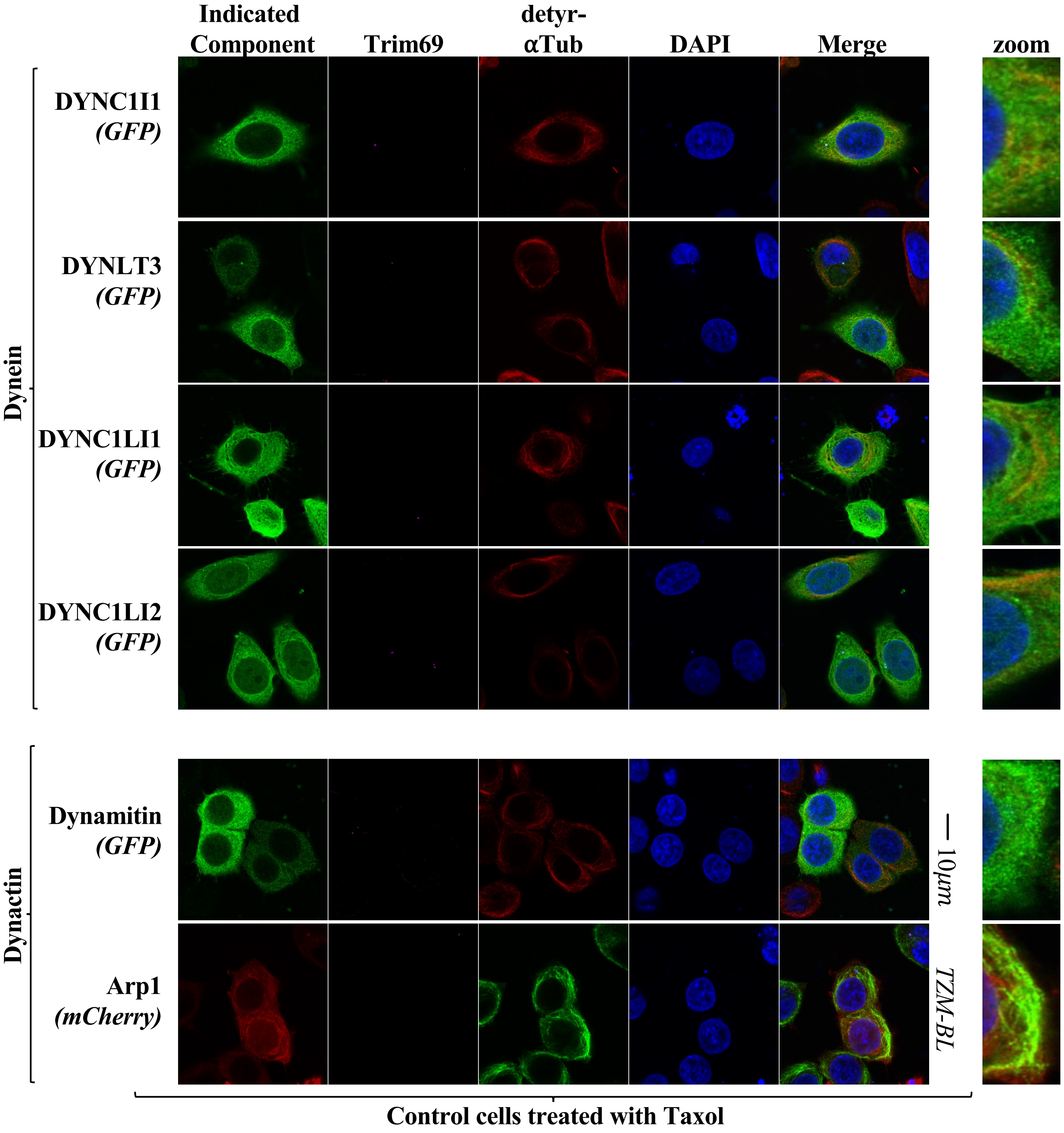

*Extended data 12*

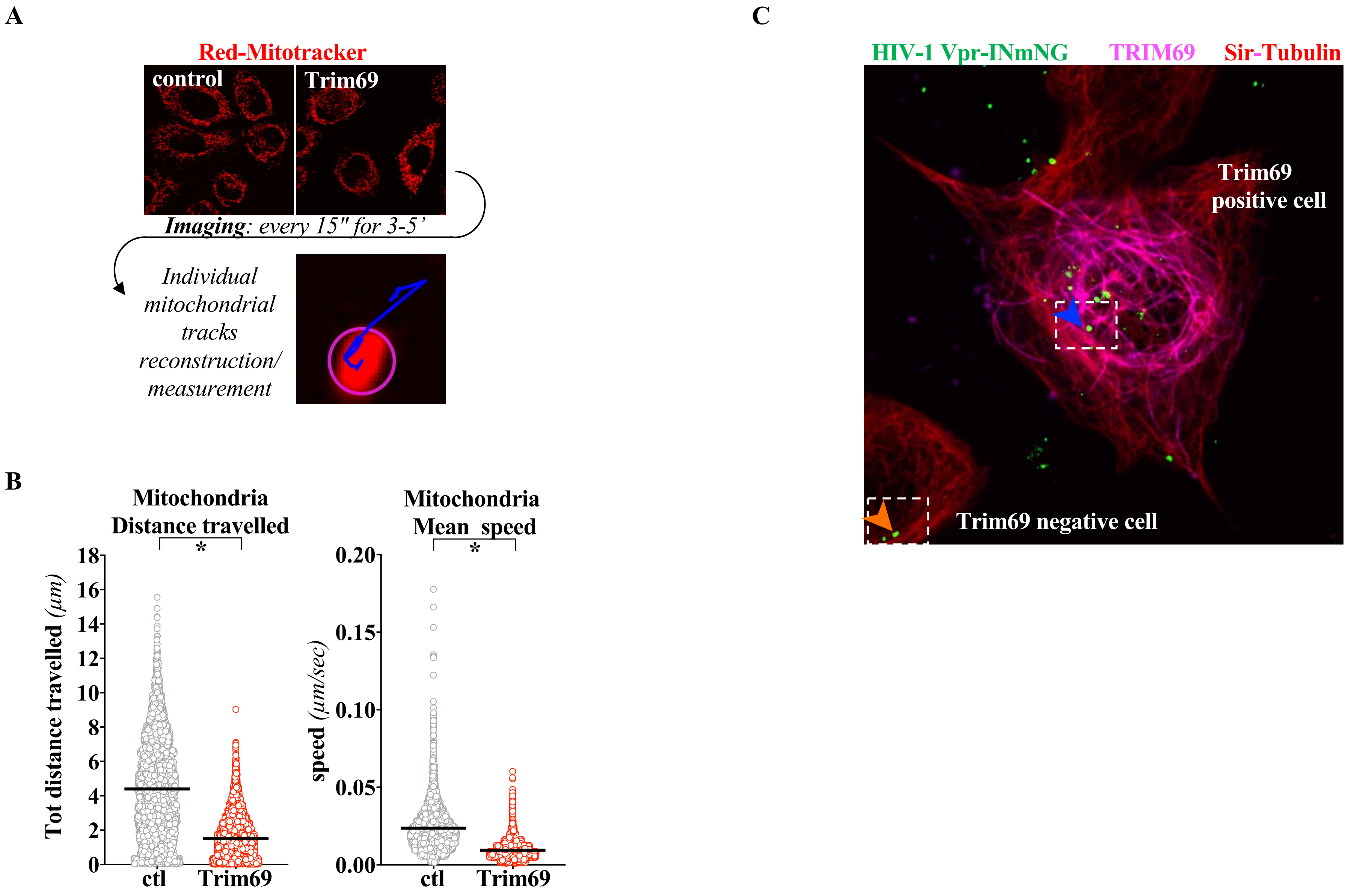
